# Proteasome gene expression is controlled by the coordinated functions of multiple transcription factors

**DOI:** 10.1101/2023.04.12.536627

**Authors:** Jennifer E. Gilda, Asrafun Nahar, Dharanibalan Kasiviswanathan, Nadav Tropp, Tamar Gilinski, Tamar Lahav, Yael Mandel-Gutfreund, Soyeon Park, Shenhav Cohen

## Abstract

Proteasome activity is crucial for cellular integrity, but how tissues adjust proteasome content in response to catabolic stimuli is uncertain. Here, we demonstrate that transcriptional coordination by multiple transcription factors is required to increase proteasome content and activate proteolysis in catabolic states. Using denervated mouse muscle as a model system for accelerated proteolysis *in vivo*, we reveal that a two-phase transcriptional program activates genes encoding proteasome subunits and assembly chaperones to boost an increase in proteasome content. Initially, gene induction is necessary to maintain basal proteasome levels, and in a more delayed phase (7-10 d after denervation) it stimulates proteasome assembly to meet cellular demand for excessive proteolysis. Intriguingly, the transcription factors PAX4 and α-PAL^NRF-1^ control the expression of proteasome among other genes in a combinatorial manner, driving cellular adaptation to muscle denervation. Consequently, PAX4 and α-PAL^NRF-1^ represent new therapeutic targets to inhibit proteolysis in catabolic diseases (e.g. type-2 diabetes, cancer).

## INTRODUCTION

Proteasome function is essential for vitality of all cells. By catalyzing the degradation of most cellular proteins (normal, unfolded, misfolded or damaged), the proteasome sustains cellular integrity, biological functions and tissue homeostasis. Conversely, impaired proteasome function is associated with protein accumulation and aggregation in age-related pathologies and neurodegenerative diseases ^1^. To withstand such diseases and compensate for the lost proteasome function, cells appear to have evolved mechanisms to increase proteasome production ^2–4^. Raising proteasome content is also important for survival during fasting, when accelerated proteolysis in muscle produces amino acids that are converted to glucose by the liver to nurture the brain. Such physiological demands are met via regulation of proteasome abundance by activation of proteasome gene expression or by post-synthetic mechanisms. Certain transcription factors have been reported to increase the expression of specific proteasome subunit genes to maintain basal levels of assembled active proteasomes ^5, 6^, or to respond to proteasome inhibition in cultured cells ^7, 8^. However, a more global response to boost proteasome content (expression and assembly) is probably necessary to meet cellular demand for excessive proteolysis in physiological catabolic states.

Coordinated induction of proteasome subunits and various ubiquitin-proteasome system (UPS) components is primarily responsible for the increased degradation of muscle proteins during atrophy ^9–11^. The resulted loss of muscle mass and strength is an inevitable sequel of many systemic catabolic states (e.g. cancer, inactivity, and malnutrition), and leads to frailty, disability, morbidity and mortality ^12^. Because proteasome subunits are induced in most types of atrophy, animal models for muscle atrophy serve as an optimal *in vivo* system in a whole organism to address critical questions related to proteasome gene expression, assembly and regulation

The proteasome is a large, multiprotein complex composed of at least 33 subunits. The 20S core contains the proteolytic sites shielded within a cylindrical chamber, which may be capped by a regulatory particle, the most common of which is the 19S regulator. Six AAA-ATPase subunits (Rpt1-6) form the base of the 19S cap, and are responsible for unfolding substrates, opening the chamber of the 20S core, and translocating substrates to the proteolytic core ^13^. The remainder of the 19S cap is composed of non-ATPase subunits (Rpn subunits), which recognize polyubiquitinated protein substrates and remove ubiquitin from them ^14^. At least five chaperones are involved in the assembly of the 20S core, namely proteasome assembly chaperones (PAC) 1-4 and proteasome maturation protein (POMP), while four chaperones are involved in the assembly of the Rpt ring, which are conserved between yeast and mammals: PAAF1/Rpn14, Nas6/gankyrin, Nas2/p27, and Hsm3/S5b ^15^. With 33 subunits required to work in concert, proteasome gene induction has to be centrally orchestrated by transcription factors. Here, we report a novel mechanism elevating proteasome production *in vivo*, which involves the coordinated functions of two transcription factors, PAX4 and α-PAL^NRF-1^.

We recently studied in mice the atrophy induced by muscle denervation, and uncovered a delayed phase in the atrophy process, which involves the induction of genes that promote proteolysis by the transcription factor PAX4. PAX4 was originally identified as a transcriptional repressor in beta islet cells during pancreatic development ^16–18^, and its potential roles in muscle (or other tissues) had been generally overlooked. We discovered that this transcription factor induces AAA-ATPase p97/VCP, the ubiquitin ligases MuRF1 and NEDD4, and the proteasome subunit PSMC2 (Rpt1) in the late stage of atrophy (10 d after denervation), and its function is crucial for degradation of ubiquitinated contractile myofibrils ^19^. This second phase of gene expression during atrophy, when myofibril breakdown is rapid, occurs long after induction of the major atrophy-related genes by the transcription factors FOXO3 ^20, 21^.

An established transcription factor for proteasome genes in cultured cells treated with proteasome inhibitors is nuclear factor, erythroid 2 like 1, known as NRF-1 (gene name *NFE2L1*) ^7, 8^. Here, we investigated another ubiquitously expressed transcription factor, nuclear respiratory factor 1, known as NRF-1 or α-PAL. Because these two distinct genes are commonly confused in the literature due to nearly identical abbreviations, we adopted the abbreviations used by Zhang: NRF-1^NFE2L1^ (Nuclear factor, erythroid 2 like 1) and α-PAL^NRF-1^ (nuclear respiratory factor 1) ^22^. ChIP sequencing data from cultured SK-N-SH human neuroblastoma cells and analyses of the Encyclopedia of DNA Elements (ENCODE) data suggested that α-PAL^NRF-1^ may regulate proteasome genes ^23, 24^; however, these data were not further verified. We show here that α-PAL^NRF-1^ is of prime importance in the induction of most proteasome genes in denervated mouse muscle *in vivo*. Its function seems to be coordinated with PAX4, representing a new mode of regulation of proteasome gene expression.

## RESULTS

### Two-phase differential expression of proteasome genes in atrophying mouse muscle *in vivo*

To understand proteasome dynamics during adaptation to a changing physiological environment, we studied denervated mouse muscles, which due to accelerated proteolysis primarily by the UPS lose ∼60% of their mass in 28 d (Fig. 1A). During the slow atrophy induced by denervation, proteolysis gradually increases over time, allowing investigation of multiple cellular phases of this debilitating process ^19^. The sciatic nerve of adult mice was transected, and the muscles were analyzed 3 – 28 d later and compared to innervated controls. Denervation led to a gradual decrease in mass of tibialis anterior (TA) (13.6% loss at 3 d and 56.3% loss at 28 d) and gastrocnemius (GA) (11.5% loss at 3 d and 65.4% loss at 28 d) skeletal muscles (Fig. 1A). This loss of mass was accompanied by a gradual decrease in muscle fiber size (cross-sectional area) (Fig. 1B and Table 1), and the absolute content of the insoluble fraction (cytoskeletal and myofibrillar proteins, which comprise 70% of muscle proteins) (Fig. 1C). These data are in line with previous findings ^11, 25, 26^.

**Figure 1.**
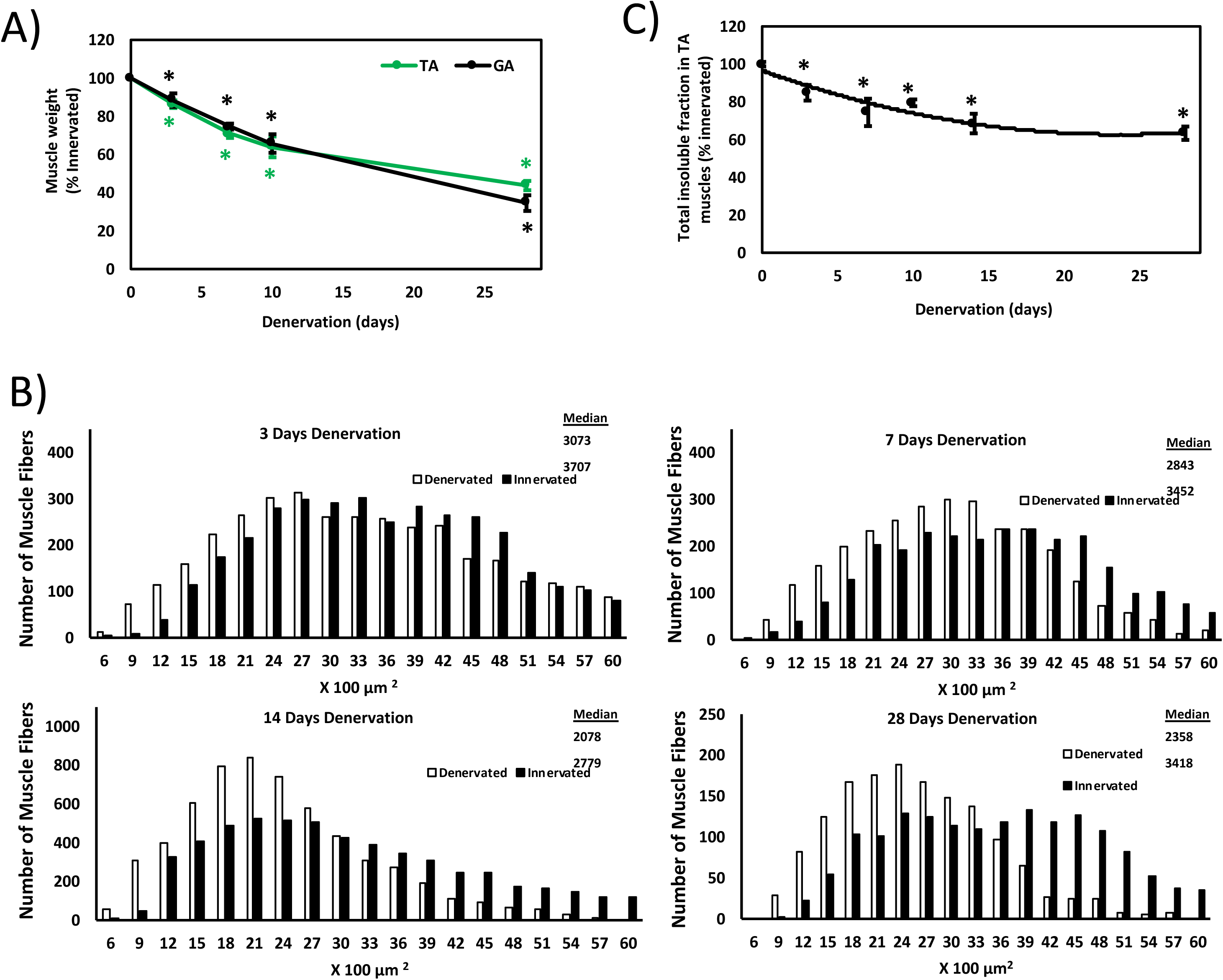
Time course of mouse muscle atrophy after denervation. The sciatic nerve of adult WT mice was transected, and tibialis anterior (TA) and gastrocnemius (GA) muscles were collected at several time points later. Muscle weight (A), muscle fiber size (B), and total amount of insoluble fraction (C) decrease after denervation. The atrophy seen over time results primarily from the accelerated degradation of muscle proteins. A) relative mean muscle weights of denervated TA and GA muscles as the percentage of innervated controls. N=5. *, p < 0.05 *vs.* innervated muscles. Error bars represent SEM. B) Measurements of cross-sectional areas of 5143 (3 d, n= 4 mice), 2887 (7 d, n= 4 mice), 5878 (14 d, n= 4 mice), 1480 (28 d, n=4 mice) fibers in denervated muscles and a similar number of fibers in innervated controls. C) Mean total content of insoluble fraction per TA muscle at different times after denervation is depicted as the percent of innervated controls. N=3. *, p < 0.05 *vs.* innervated muscles. Error bars represent SEM.

**Table 1.**
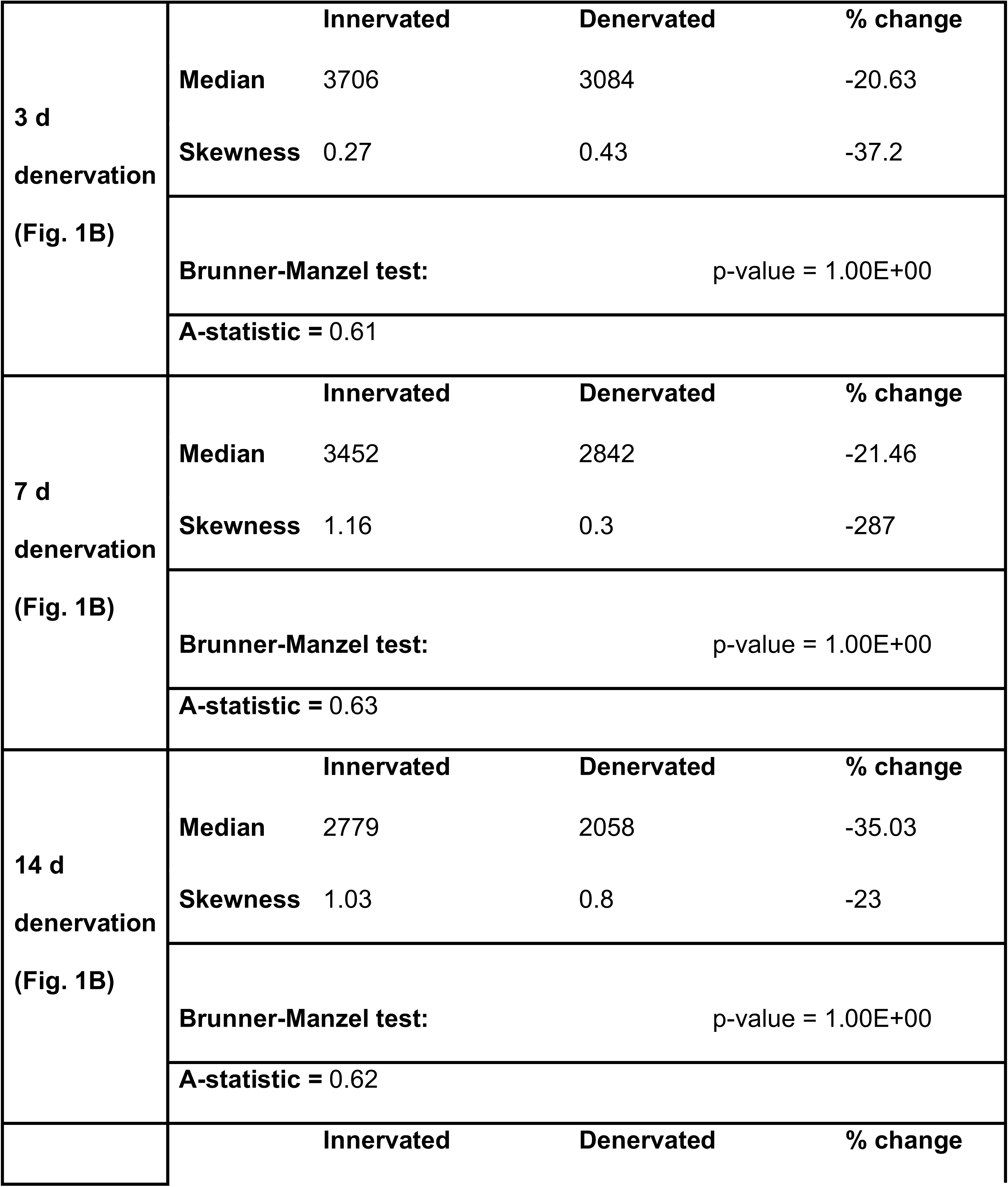
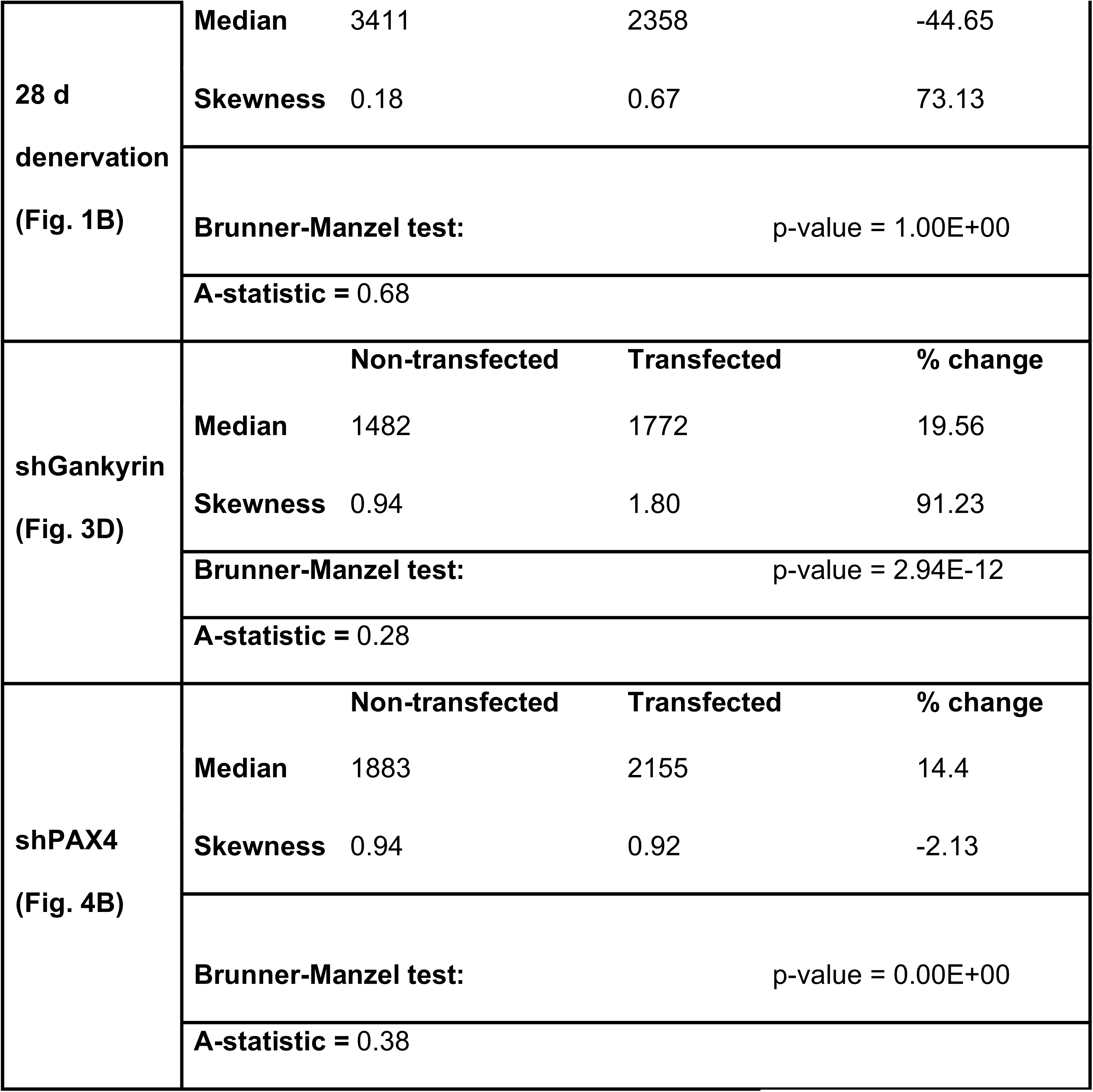
Statistical analysis of fiber size measurements. .Summary statistics of fiber size analyses presented in Figs. 1B, 3D and 4B based on our recent methodology paper ^52^. With regard to A-statistics, if 0≤A<0.5 then dataset1 (non-transfected) is stochastically less than dataset2 (transfected). If 0.5≤A<1 then dataset1 (innervated) is stochastically greater than dataset2 (denervated). The A-statistics is a direct measure of the fiber size effect ^52^, and it shows beneficial effect on cell size by the specific shRNAs. Such an effect can be simply missed by traditional measurements of median, average, and Student’s *t*-test.

Our prior investigations concluded that atrophy is a two-phase process involving an early (3-7 d post-denervation) loss of the desmin cytoskeleton, followed by a more delayed (at 10 – 14 d) increase in myofibril ubiquitylation and degradation by the UPS^19, 26^. While the rate of muscle loss is highest early in atrophy (3d after denervation) ^11^, overall protein degradation appears to be highest in the late phase of atrophy ^26–28^. To determine how the expression of UPS genes is coordinated within these two phases of protein degradation, we analyzed atrophying muscles at different times after denervation by RT-PCR and compared these data to innervated controls. Expression of proteasome subunit genes was analyzed according to subunit type along with their corresponding chaperones. Prior investigations on cultured cells or yeast showed that all proteasome genes rise at the same time ^29^. However, when investigated in atrophying adult mouse skeletal muscles *in vivo*, we surprisingly show distinct dynamics of proteasome gene expression during physiological adaptation to catabolic cues (e.g., muscle denervation). Gene expression of PSMC1 (Rpt2), PSMC4 (Rpt3), and PSMC5 (Rpt6) increased by 1.5-2.5 fold on average by 3 d after denervation, and remained fairly steady until 28 d (Fig. 2A). The induction of PSMC4 (Rpt3) and PSMC5 (Rpt6) at an early phase in this debilitating process is in line with prior reports demonstrating the module formation of PSMC4 (Rpt3)-PSMC5 (Rpt6) and associated chaperones as the rate limiting step for overall proteasome assembly ^30–32^. The gene expression pattern of PSMC2 (Rpt1), PSMC6 (Rpt4), and PSMC3 (Rpt5) showed more dynamic two-phase behavior, with higher overall induction and peaks in expression at 3-7 and 14 d after denervation (Fig. 2B).

**Figure 2.**
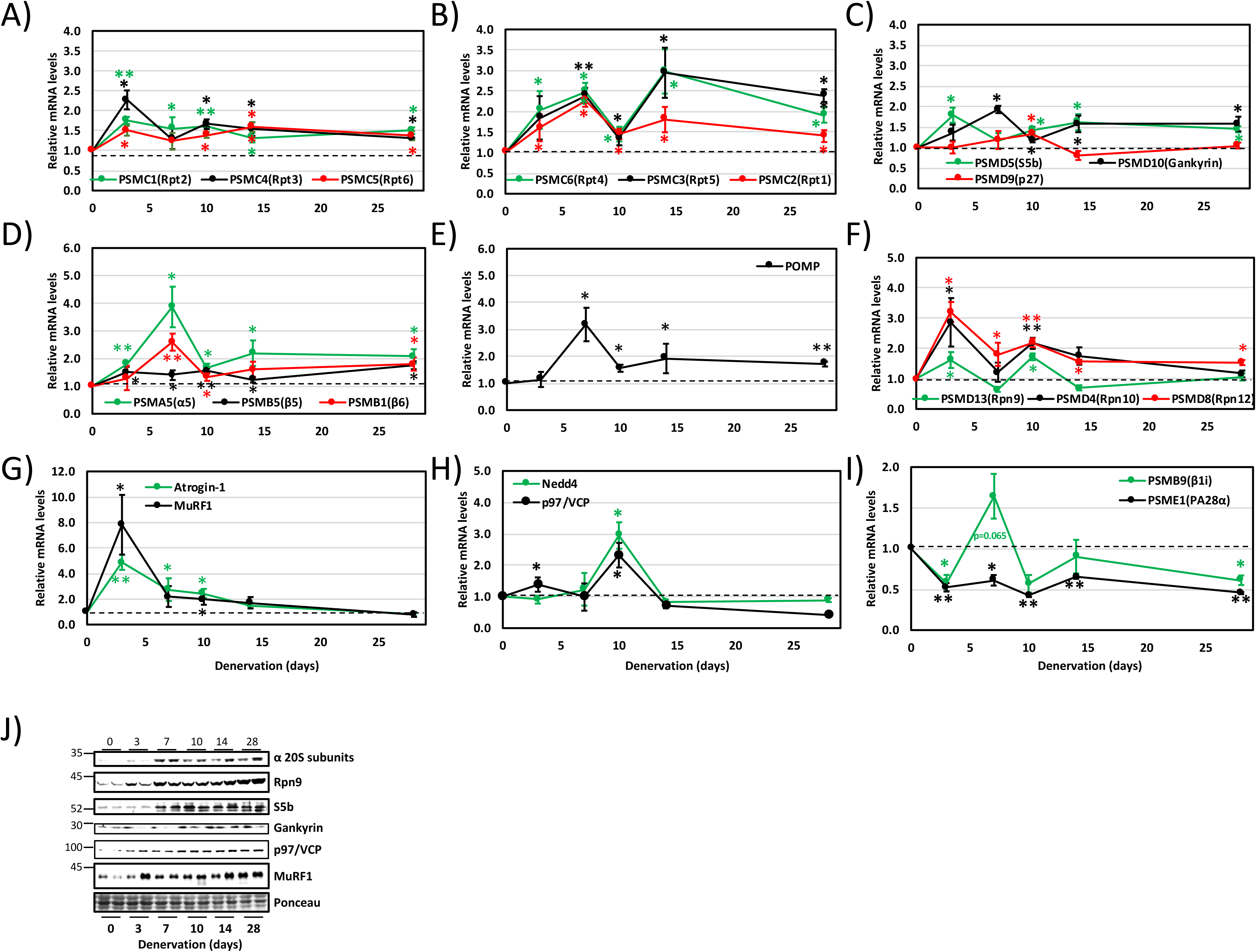
Two-phase differential expression of proteasome genes in atrophying mouse muscle *in vivo*. A – I) Time course of induction of selected proteasome and chaperone genes was determined by RT-PCR analysis of mRNA preparations from TA muscles at 3, 7, 10, 14, and 28 d after denervation. N=7. *, P < 0.05; ** P < 0.001 *vs.* innervated muscles. Error bars represent SEM. J) Increased protein levels were confirmed for representative genes by analysis of soluble fractions from innervated and denervated muscles by SDS-PAGE and immunoblotting.

The Rpt subunits are first assembled into dimers before incorporation into the hexameric ring that forms the base of the 19S cap. PSMC2 (Rpt1) and PSMC1 (Rpt2) are assembled into a heterodimer by the chaperone PSMD5 (S5b); PSMC4 (Rpt3) and PSMC5 (Rpt6) by the chaperone gankyrin, and PSMC6 (Rpt4) and PSMC3 (Rpt5) by p27^33^. The genes encoding PSMD5 (S5b), PSMD10 (gankyrin), and PSMD9 (p27) were all induced by 10 d after denervation (Fig. 2C). These data suggest that chaperones that are present in denervated muscle at 3 d after denervation should be sufficient to maintain basal proteasome biogenesis, and basal proteasome levels are sufficient to carry out the muscle loss that occurs at this early phase of the atrophy process (3-7 d after denervation). Then, chaperone levels are elevated at 7 d, most likely to induce proteasome assembly, just when myofibril disassembly and destruction is accelerated (10-14 d after denervation) ^19, 26, 34^.

To test whether other proteasome subunits are induced in this delayed phase of atrophy, we examined the expression of 20S subunits PSMA5 (α5), PSMB5 (β5), and PSMB1 (β6), which were all elevated in denervated muscle by day 7 (Fig. 2D). Gene expression of the 20S chaperone POMP also peaked at 7 d after denervation (Fig. 2E), most likely to boost proteasome biogenesis just before protein degradation is accelerated ^19, 26^. In addition, induction of Rpn subunits [PSMD13 (Rpn9), PSMD4 (Rpn10), and PSMD8 (Rpn12)] initially peaked at 3 d after denervation followed by a second wave of induction at 10 d (Fig. 2F).

Expression of most genes returned toward baseline at 14 d after denervation and showed a sustained low mRNA levels until 28 d (Fig. 2A-F). Accordingly, RNA sequencing (RNA-Seq) analysis of TA muscles at 14 d after denervation indicated that expression of most proteasome genes is low at 14 d (Fig. S1). Relatively few changes were seen in proteasome gene transcript levels [9 subunits were significantly induced in denervated compared to innervated controls, including PSMA7 (α4), PSMA5 (α5), PSMB5 (β5), PSMB4 (β7), PSMB3 (β3), PSMB1 (β6), PSMD8 (Rpn12), PSMD13 (Rpn9) and PSMD4 (Rpn10)(Fig. S1).

Further studies determined the expression patterns of key UPS components that are known to promote proteolysis in muscle atrophy. The ubiquitin ligases MuRF1 and atrogin-1, which are required for atrophy, peaked at 3 d and then returned to basal levels (^11, 21^) (Fig. 2G). Distinctly, gene expression of the atrophy-related genes Nedd4 and p97/VCP peaked at a more delayed phase (10 d) (Fig. 2H) as we and others had demonstrated before ^19, 35^. Together, these findings systemically extend our prior studies on PSMC2 (Rpt1) ^19^ and indicate early (certain Rpt and Rpn subunits, MuRF1, atrogin-1) and delayed (20S, certain Rpt and Rpn subunits, Nedd4, and p97/VCP) phases of induction of proteasome genes and UPS components during the atrophy process.

It is noteworthy that the transcript levels of the alternate regulatory particle PSME1 (PA28α) were decreased by ∼50% at all times after denervation (Fig. 2I), and the expression of the inducible catalytic subunit of the immunoproteasome, PSMB9 (β1i) was decreased or unchanged following denervation (Fig. 2I), when 26S chaperones (Fig. 2C and E) and proteasome assembly (see below, Fig. 3) were increased, suggesting the specific induction of biogenesis of the major constitutive 26S proteasomes holoemzyme in atrophying mouse muscles.

**Figure 3.**
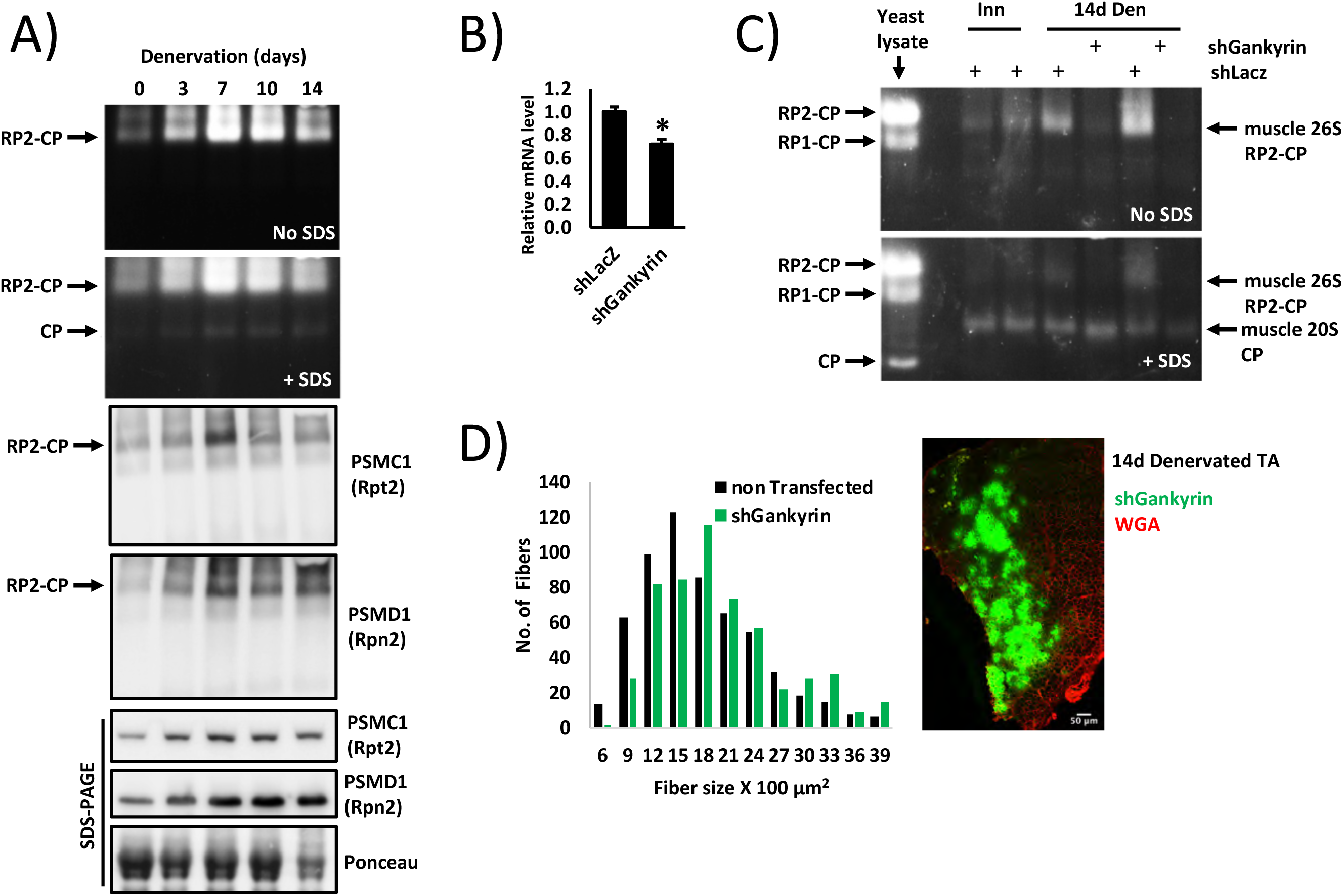
Increased proteasome assembly is a delayed response to muscle denervation. A) Innervated and denervated muscle homogenates were analyzed by native polyacrylamide gel electrophoresis, and in-gel LLVY-AMC (β5 proteasome substrate) cleavage was visualized by UV light. Top two panels: 30S activity is shown in the absence of SDS, and 20S activity was detected by the addition of SDS. Lower panels: Western blotting for PSMC1(Rpt2) and PSMD1 (Rpn2) in native and denaturing gels. B) shGankyrin downregulates gankyrin in NIH-3T3 cells. mRNA preparations from transfected cells were analyzed by RT-PCR and specific primers to gankyrin. N=3. *, P < 0.05 *vs.* shLacz. Error bars represent SEM. C) Gankyrin is required for proteasome assembly in atrophying mouse muscles. Innervated and denervated TA muscles expressing shGankyrin or scrambled shLacz (control) were analyzed by native gel and in-gel LLVY-AMC cleavage assay D) Downregulation of gankyrin in denervated muscles reduces fiber atrophy. Measurements of cross-sectional areas of 590 fibers expressing shGankyrin and adjacent 590 non-transfected fibers. N= 5 mice. Representative image of electroporated muscle is shown. Scale bar: 50 μm.

Western Blot analysis of representative genes indicated that this increase in mRNA levels largely correlated with protein levels, which generally increased at 7 d and remained elevated till 28 d (Fig. 2J). Although Rpn9 gene expression pattern followed two waves of induction, its protein levels increased substantially only by 7 d (Fig. 2J), just before muscle protein degradation is accelerated. Moreover, MuRF1 mRNA peaked at 3 d and then returned to basal levels, but its protein levels remained elevated until 28 d (Fig. 2J). In addition, p97/VCP peaked at a more delayed phase (10 d) while the protein levels of p97/VCP showed a moderate, steady increase (Fig. 2J).

### Increased proteasome assembly is a delayed response to muscle denervation

Since PSMD9 (p27), the chaperone for PSMC6 (Rpt4) and PSMC3 (Rpt5), is induced later (10 d, Fig. 2C), the increase in proteasome biogenesis was expected in a more delayed phase, when new assembly modules are formed via gene expression. Consequently, chaperones and assembly modules that are present in muscle between 3-7 d after denervation are probably sufficient to meet cellular demand. To determine whether the observed increase in proteasome gene and protein expression is coordinated with increased proteasome assembly, we assayed proteasome activity in innervated and denervated muscles by native gel electrophoresis and in-gel AMC-LLVY cleavage, which measures the β5 proteolytic activity of the proteasome. Because most changes in gene expression were observed from 3 – 14 d, homogenates of denervated muscles at 3 – 14 d after denervation and innervated controls were analyzed. As shown in Fig. 3A, total activity of 30S (doubly capped) proteasomes was increased already by 3 d, but was dramatically elevated at 7-10 d (Fig. 3A upper panel), when the expression of most 26S subunits and chaperones is elevated (Fig. 1C-E), and just before myofibril breakdown is accelerated ^26^. By contrast, the overall changes in activity of the 20S core particle (CP) were much lower, suggesting that the excessive proteolysis in atrophying muscle is carried out primarily by the singly (26S, RP1-CP) or doubly (30S, RP2-CP) capped proteasomes (Fig. 3A). Western blotting of native and SDS gels for the 19S subunits PSMC1 (Rpt2) and PSMD1 (Rpn2) confirmed that the increase in activity is due, at least in part, to an increase in abundance of 30S proteasomes (Fig. 3A).

The induction of the proteasome chaperones POMP, PSMD5 (S5b), PSMD10 (gankyrin), and PSMD9 (p27) in denervated muscles (Fig. 2C and E) is probably required to boost proteasome biogenesis in atrophying muscles. To test this idea, we downregulated gankyrin by the electroporation of a specific shRNA (shGankyrin) into TA muscles, which reduced gankyrin mRNA levels below the levels in shLacz expressing controls (Fig. 3B). This *in vivo* electroporation technique is a powerful method for determining transient gene effects on proteolysis and cell size within days ^36, 37^. To assess the cumulative effects of chaperone downregulation, we analyzed transfected muscles at 14 d after denervation (Fig. 3C). Analysis of innervated and 14 day denervated muscles expressing shGankyrin or control (shLacz) by in-gel activity assay indicated that downregulation of the chaperone gankyrin was sufficient to dramatically reduce proteasome assembly in denervated muscles, below baseline levels of assembled 30S proteasomes in innervated controls (Fig. 3C). This decrease in assembled proteasomes dramatically attenuated protein breakdown and atrophy because the cross-sectional area (CSA) of 14 d denervated muscle fibers expressing shGankyrin (also express GFP) was significantly greater than adjacent non-transfected fibers (Fig. 3D). Thus, a second wave of gene induction (7-10 d after denervation), when most proteasome genes and assembly chaperons (e.g., gankyrin) are elevated, seems necessary to boost proteasome biogenesis and satisfy the physiological demand for accelerated proteolysis.

### PAX4 induces proteasome subunit genes *in vivo*

We previously showed that PAX4 is required for the induction of the proteasome subunit PSMC2 (Rpt1) and the atrophy-related genes MuRF1, p97/VCP, and Nedd4 at 10 d after denervation ^19^. To determine if PAX4 controls the expression of other proteasome subunits and/or chaperones, we searched the promoter regions of these genes for potential PAX4 binding sites using the FIMO (Find Individual Motif Occurrences) algorithm ^38^, using the human PAX4 motif (MA0068.2). We found that 13 proteasome genes contained potential binding sites for the transcription factor PAX4 (Table 2), suggesting that PAX4 was likely stimulating their induction in denervated muscle. To clarify the role of PAX4 *in vivo*, we transiently suppressed its expression by electroporation into denervated mouse TA of shRNA plasmid (shPAX4, validated in ^19^). The expression of all 13 proteasome genes that were analyzed as well as of the ubiquitin ligases MuRF1 and atrogin-1 significantly increased at 10 day after denervation (Fig. 4A); however, the downregulation of PAX4 with shPAX4 resulted in a marked decreased in the expression of PSMA5 (α5), PSMC2 (Rpt1), PSMC1 (Rpt2), PSMC3 (Rpt5), MuRF1, and atrogin-1, indicating that PAX4 is required for their induction (Fig. 4A). Thus, on denervation, PAX4 is required for the induction of genes that promote ubiquitination and proteolysis. Accordingly, downregulation of PAX4 significantly attenuated muscle fiber atrophy at 10 d after denervation (Fig. 4B and Table 1), a beneficial effect on muscle that results from reduced proteolysis ^36^.

**Figure 4.**
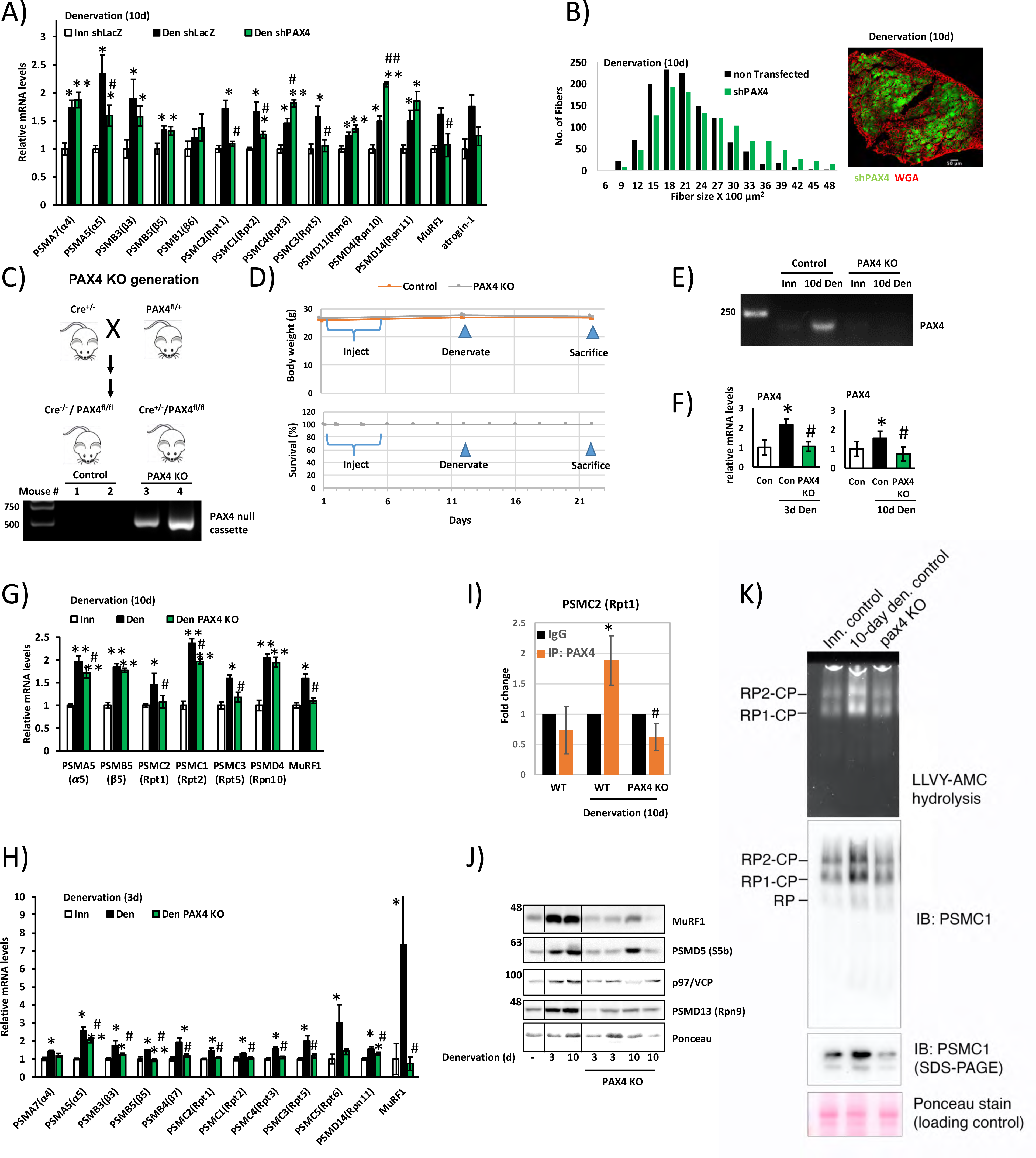
PAX4 induces proteasome genes *in vivo*. A) Proteasome gene expression was measured by RT-PCR analysis of mRNA preparations from innervated and 10 day denervated muscles expressing shPAX4 or shLacz control. N=5. *, P < 0.05 *vs.* innervated shLacz; **, P < 0.001 *vs.* innervated shLacz # P < 0.05 *vs.* denervated shLacz; ## P < 0.001 *vs.* denervated shLacz. Error bars represent SEM. B) PAX4 downregulation with shPAX4 reduces fiber atrophy on denervation. Measurement of cross-sectional areas of 1178 fibers from 10 d denervated fibers expressing shPAX4 *vs.* 1178 adjacent non-transfected fibers. N= 3 mice. Right: representative image of transfected muscle. Scale bar: 50 μm. C) Top: Strategy for generating PAX4 conditional KO mice. Bottom: DNA was isolated from tail snippings and analyzed by PCR genotyping. Following tamoxifen injections of Cre^+/-^/PAX4^fl/fl^ mice, removal of PAX4 from the genome was verified using the P1/P3 primers (Supplemental Table S3). D) Body weight (g) and survival (%) of PAX4 KO and WT mice are presented, following tamoxifen injections and muscle denervation. E-F) PAX4 mRNA increases on denervation of WT mouse muscle, but is absent in TA muscles from PAX4 KO mice. Mice were injected with tamoxifen and their muscles denervated. PCR (E) or RT-PCR (F) using primers for PAX4 was performed on mRNA preparations from innervated and denrvated muscles. G-H) Proteasome gene expression was measured by RT-PCR analysis of mRNA preparations from innervated and 10 (G) or 3 (H) days denervated muscles from WT and PAX4 KO mice. N=5. *, P < 0.05 *vs.* innervated shLacz; ** P < 0.001 *vs.* innervated shLacz; # P < 0.05 *vs.* denervated shLacz. Error bars represent SEM. I) PAX4 binds the promoter region of *PSMC2* gene. ChIP was performed on innervated and denervated (10 d) muscles from WT or PAX4 KO mice using PAX4 antibody or non-specific IgG control, and primers for *PSMC2* gene. Data is plotted as fold change relative to IgG control. N=3. *, P < 0.05 *vs.* innervated WT; # P < 0.05 *vs.* denervated WT. Error bars represent SEM. J) Correlative reduction in protein levels was confirmed for representative genes by analysis of soluble fractions from innervated and denervated muscles from WT or PAX4 KO mice by SDS-PAGE and immunoblotting. K) The content of active assembeled proteasomes increase in denervated muscles, but not in msucles lacking PAX4. Measurement of proteasome content by native gels and immunoblotting, and proteasome peptidase activity by LLVY-AMC cleavage.

**Table 2.** Potential PAX4 binding sites in mouse proteasome genes. Predicted PAX4- and α-PAL^NRF-1^-binding sites in the promoter regions of the mouse genes for proteasome subunits, and their distances from TSS.

| TF | gene_name1 | gene_name | peak_start | p_value | TF |
| --- | --- | --- | --- | --- | --- |
| PAX4 | alpha5(PSMA5) | PSMA5 | chr3:108256482 | 0.000338 | Nrf1 |
| PAX4 | alpha5(PSMA5) | PSMA5 | chr3:108256647 | 0.000426 |  |
| PAX4 | alpha4(PSMA7) | PSMA7 | chr2:180042999 | 0.000386 | Nrf1 |
| PAX4 | alpha4(PSMA7) | PSMA7 | chr2:180046232 | 0.000407 | Nrf1 |
| PAX4 | beta6(PSMB1) | PSMB1 | chr17:15502502 | 0.000942 | Nrf1 |
| PAX4 | beta6(PSMB1) | PSMB1 | chr17:15503069 | 0.000669 | Nrf1 |
| PAX4 | beta3(PSMB3) | PSMB3 | chr11:97700415 | 0.000457 | Nrf1 |
| PAX4 | beta3(PSMB3) | PSMB3 | chr11:97703007 | 0.000312 | Nrf1 |
|  |  |  |  |  | Nrf1 |
| PAX4 | beta5(PSMB5) | PSMB5 | chr14:54622572 | 0.000386 | Nrf1 |
| PAX4 | beta5(PSMB5) | PSMB5 | chr14:54622784 | 0.000457 |  |
| PAX4 | Rpt2(PSMC1) | PSMC1 | chr12:100109305 | 4.83e-05 | Nrf1 |
| PAX4 | Rpt2(PSMC1) | PSMC1 | chr12:100109413 | 4.83e-05 |  |
| PAX4 | Rpt1(PSMC2) | PSMC2 | chr5:21780915 | 4.83e-05 |  |
| PAX4 | Rpt1(PSMC2) | PSMC2 | chr5:21783713 | 0.000337 |  |
| PAX4 | Rpt5(PSMC3) | PSMC3 | chr2:91051955 | 0.000188 | Nrf1 |
| PAX4 | Rpt5(PSMC3) | PSMC3 | chr2:91052805 | 0.000163 |  |
| PAX4 | Rpt3(PSMC4) | PSMC4 | chr7:28050111 | 0.000141 | Nrf1 |
| PAX4 | Rpt3(PSMC4) | PSMC4 | chr7:28054285 | 0.000337 | Nrf1 |
|  |  |  |  |  | Nrf1 |
|  |  |  |  |  | Nrf1 |
|  |  |  |  |  | Nrf1 |
|  |  |  |  |  | Nrf1 |
|  |  |  |  |  | Nrf1 |
|  |  |  |  |  | Nrf1 |
| PAX4 | Rpt6(PSMC5) | PSMC5 | chr11:106253444 | 0.000141 | Nrf1 |
| PAX4 | Rpt6(PSMC5) | PSMC5 | chr11:106254031 | 0.000407 | Nrf1 |
|  |  |  |  |  | Nrf1 |
| PAX4 | Rpn6(PSMD11) | PSMD11 | chr11:80427775 | 0.000235 | Nrf1 |
| PAX4 | Rpn6(PSMD11) | PSMD11 | chr11:80427479 | 0.000444 | Nrf1 |
|  |  |  |  |  | Nrf1 |
|  |  |  |  |  | Nrf1 |
| PAX4 | Rpn11(PSMD14) | PSMD14 | chr2:61710225 | 0.000312 | Nrf1 |
| PAX4 | Rpn11(PSMD14) | PSMD14 | chr2:61709754 | 0.000337 | Nrf1 |
|  |  |  |  |  | Nrf1 |
| PAX4 | Rpn10(PSMD4) | PSMD4 | chr3:95045797 | 4.83e-05 | Nrf1 |
| PAX4 | Rpn10(PSMD4) | PSMD4 | chr3:95046650 | 0.000235 |  |
| PAX4 | VCP(p97) | p97 | chr4:43000801 | 9.08e-05 |  |
|  |  |  |  |  | Nrf1 |
|  |  |  |  |  | Nrf1 |
|  |  |  |  |  | Nrf1 |

| gene_name1 | gene_name | peak_start | p_value |
| --- | --- | --- | --- |
| alpha5(PSMA5) | PSMA5 | chr3:108256070 | 5.99e-05 |
| alpha4(PSMA7) | PSMA7 | chr2:180043041 | 5.11e-05 |
| alpha4(PSMA7) | PSMA7 | chr2:180043272 | 6.25e-05 |
| beta6(PSMB1) | PSMB1 | chr17:15499203 | 5.4e-05 |
| beta6(PSMB1) | PSMB1 | chr17:15499204 | 1.61e-05 |
| beta3(PSMB3) | PSMB3 | chr11:97700106 | 8.2e-05 |
| beta3(PSMB3) | PSMB3 | chr11:97700482 | 0.000112 |
| beta3(PSMB3) | PSMB3 | chr11:97700569 | 0.000117 |
| beta5(PSMB5) | PSMB5 | chr14:54618709 | 1.38e-05 |
| Rpt2(PSMC1) | PSMC1 | chr12:100106560 | 2.75e-06 |
| Rpt5(PSMC3) | PSMC3 | chr2:91053800 | 9.59e-05 |
| Rpt3(PSMC4) | PSMC4 | chr7:28050967 | 2.85e-05 |
| Rpt3(PSMC4) | PSMC4 | chr7:28050964 | 6.71e-05 |
| Rpt3(PSMC4) | PSMC4 | chr7:28050940 | 5.42e-06 |
| Rpt3(PSMC4) | PSMC4 | chr7:28050966 | 8.89e-06 |
| Rpt3(PSMC4) | PSMC4 | chr7:28050968 | 6.79e-06 |
| Rpt3(PSMC4) | PSMC4 | chr7:28050969 | 1.43e-05 |
| Rpt3(PSMC4) | PSMC4 | chr7:28050939 | 1.09e-07 |
| Rpt6(PSMC5) | PSMC5 | chr11:106256133 | 5.83e-07 |
| Rpt6(PSMC5) | PSMC5 | chr11:106256134 | 3.2e-06 |
| Rpt6(PSMC5) | PSMC5 | chr11:106256115 | 1.18e-05 |
| Rpn6(PSMD11) | PSMD11 | chr11:80426184 | 0.000328 |
| Rpn6(PSMD11) | PSMD11 | chr11:80425905 | 0.000761 |
| Rpn9(PSMD13) | PSMD13 | chr7:140881656 | 0.000134 |
| Rpn9(PSMD13) | PSMD13 | chr7:140881941 | 0.000205 |
| Rpn11(PSMD14) | PSMD14 | chr2:61711612 | 1.6e-07 |
| Rpn11(PSMD14) | PSMD14 | chr2:61711611 | 6.12e-06 |
| Rpn11(PSMD14) | PSMD14 | chr2:61711605 | 9.05e-05 |
| Rpn10(PSMD4) | PSMD4 | chr3:95044905 | 3.66e-05 |
| MuRF1(TRIM63) | TRIM63 | chr4:134310493 | 8.61e-05 |
| atrogin-1(FBXO32) | FBXO32 | chr15:58215256 | 5.57E-05 |
| Nedd4 | Nedd4 | chr9:72662268 | 3.22E-05 |

Our findings indicate a novel regulation of porteasome content, which evolved to allow cellls respond to a catabolic stimulus. To determine whether PAX4 is in fact essential for the increased proteasome abundance on denervation *in vivo*, we generated an inducible PAX4 knockout (KO) mouse. PAX4 plays a crucial role in the differentiation of pancreatic islets during development, and mice lacking PAX4 die within three to four days of birth due to severe diabetes, requiring the generation of inducible PAX4 KO animals ^39, 40^. For this purpose, PAX4^floxed^ (PAX4^fl^) mice were crossed with whole body Cre^+^ mice to generate homozygous floxed PAX4 mice that were positive for Cre (Fig. 4C). Cre^+/fl^ PAX4 adult mice were then injected with tamoxifen to induce Cre expression and PAX4 KO ^41^, and recombination of the lox sites in the PAX4 gene was verified by PCR analysis (Fig. 4C). These mice showed no abnormalities and developed similarly to wild type (WT) littermates (PAX4^fl/fl^). They were used for experiments at 3-4 months of age, when their body weight was 26-30g. For age-matched PAX4 KO and controls, there was no difference in body weight or survival before or after tamoxifen injection (Fig. 4D). In agreement with shPAX4 data, in muscle fibers from PAX4 null mice atrophy was attenuated on denervation (10 d) compared with denervated muscles from WT animals (Fig. S2 and Table S1).

We have found by experience that PAX4 expression is very low in muscle, and therefore difficult to detect by standard PCR. To enhance sensitivity of the detection method, we performed two PCR reactions using nested primers. cDNA from innervated and 10d denervated muscles was amplified using primers for PAX4, and this product was then further amplified using primers that were nested inside the first set of primers. We found that in innervated muscle from mice that express PAX4 normally, PAX4 levels were barely detectable, while in denervated muscle, PAX4 mRNA levels dramatically increased (Fig. 4E), indicating that PAX4 is induced in atrophying denervated muscles. By contrast, PAX4 mRNA was absent in innervated and 10d denervated muscles from PAX4 null mice, indicating a successful knockout of PAX4 gene (Fig. 4E). To quantitatively measure PAX4 expression, we performed RT-PCR on mRNA preparations from innervated and 3d and 10d denervated muscles using specific primers for PAX4 (Table S2). We found that PAX4 gene expression was elevated at 3d and 10d after denervation (2.2-fold and 1.6-fold, respectively) (Fig. 4F), but not in muscles from PAX4 KO mice.

Similar to our observations above using shPAX4 and an *in vivo* electroporation (Fig. 4A), the deficiency of PAX4 in muscles (10 d) from PAX4 KO mice blocked the induction of various proteasome genes as well as of MuRF1 at 10 d after denervation (Fig. 4G). Because PAX4 is induced at 3 d after denervation (Fig. 4F), we determined whether it is essential for proteasome gene induction also at this early stage. Of the 10 proteasome subunit genes we tested, the induction of eight (including α, β, Rpt, and Rpn subunits) as well as of MuRF1 was blocked in 3 d denervated muscles from PAX4 null mice (Fig. 4H), indicating that PAX4 promotes proteasome gene expression also in the early phase of atrophy. To validate these effects of PAX4 deficiency on proteasome gene expression and confirm that PAX4 binds the promoter regions of these genes, we focused on a representative proteasome gene, PSMC2 (Rpt1), and performed chromatin immunoprecipitation (ChIP) from innervated and denervated muscles from WT and PAX4 KO mice using a PAX4 specific antibody or a non-specific IgG as control. Analysis by RT-PCR of the immunoprecipitated DNA using specific primers for a predicted PAX4 motif within mouse PSMC2 (Rpt1) promoter ^19^ confirmed enhanced binding of PAX4 to this gene on denervation compared with innervated controls (Fig. 4I), but not in control samples containing a non-specific IgG or in muscles lacking PAX4 (i.e. from PAX4 KO mice) (Fig. 4I). Thus, PAX4 induces various proteasome genes and UPS components that promote the accelerated proteolysis and atrophy *in vivo*.

Consequently, we tested more directly if this PAX4-dependent induction of proteasome genes triggers the increase in the cellular content of assembled proteasomes during adaptation to catabolic conditions *in vivo*. However, it was initially important to determine whether the changes observed in gene expression were reflected in the protein products. We focused our attention on representative proteasome genes and UPS components, MuRF1, PSMD5, p97/VCP, PSMD13, which were induced in atrophying muscles (Fig. 2). Analysis of muscle homogenates by SDS-PAGE and immunoblotting using specific antibodies indicated that protein abundance correlated well with the transcript levels (Fig. 4J). Moreover, the dramatic increase in the protein levels of MuRF1, PSMD5 (S5b), p97/VCP, and PSMD13 (Rpn9) caused by muscle denervation was completely abrogated in denervated muscles lacking PAX4 (i.e., from PAX4 KO mice) (Fig. 4J).

Measurement of proteasome content by native gels and immunoblotting, and proteasome peptidase activity by LLVY-AMC cleavage, confirmed an increase in active assembled proteasomes in mouse muscle at 10 d after denervation, but not in atrophying muscles lacking PAX4 (Fig. 4K). This elevated proteasomal content must increase the cell’s capacity for proteolysis and therfore is critical in the process of muscle wasting. The inhibition of proteasome production on PAX4 deficiency should account for the resulting attenuation in muscle wasting (Fig. 4B).

### α-PAL^NRF-1^ is a novel transcription factor required for proteasome gene induction *in vivo*

Because PAX4 regulated some but not all proteasome genes in denervation-induced atrophy, we sought to investigate the roles of additional transcription factors. FOXO3 is a transcription factor known to control the expression of multiple atrophy-related genes in the early phase of atrophy ^42^. To examine its role in promoting proteasome gene induction during the late phase of atrophy, which was not previously investigated, we electroporated TA muscles with a plasmid encoding a dominant negative inhibitor of FOXO3 (FOXO3ΔC) and analyzed the effects on proteasome gene induction by RT-PCR. The FOXO3ΔC construct lacks the C-terminal transactivation domain, which is required for the transactivation of its target genes ^43^. At 3 d after denervation, FOXO3ΔC blocked the induction of the ubiquitin ligase MuRF1, consistent with prior reports ^20^, but not of proteasome subunits (Fig. 5A)(as shown before ^42^). Interestingly, in muscles denervated for 10 d, FOXO3 inhibition by the expression of FOXO3ΔC blocked the induction of only two proteasome subunit genes, PSMD11 (Rpn6) and PSMD13 (Rpn9), as well as of MuRF1 and atrogin-1 (Fig. 5B). Hence, FOXO3 appears to promotes muscle loss primarily by regulating other atrophy-related targets besides proteasome subunits.

**Figure 5.**
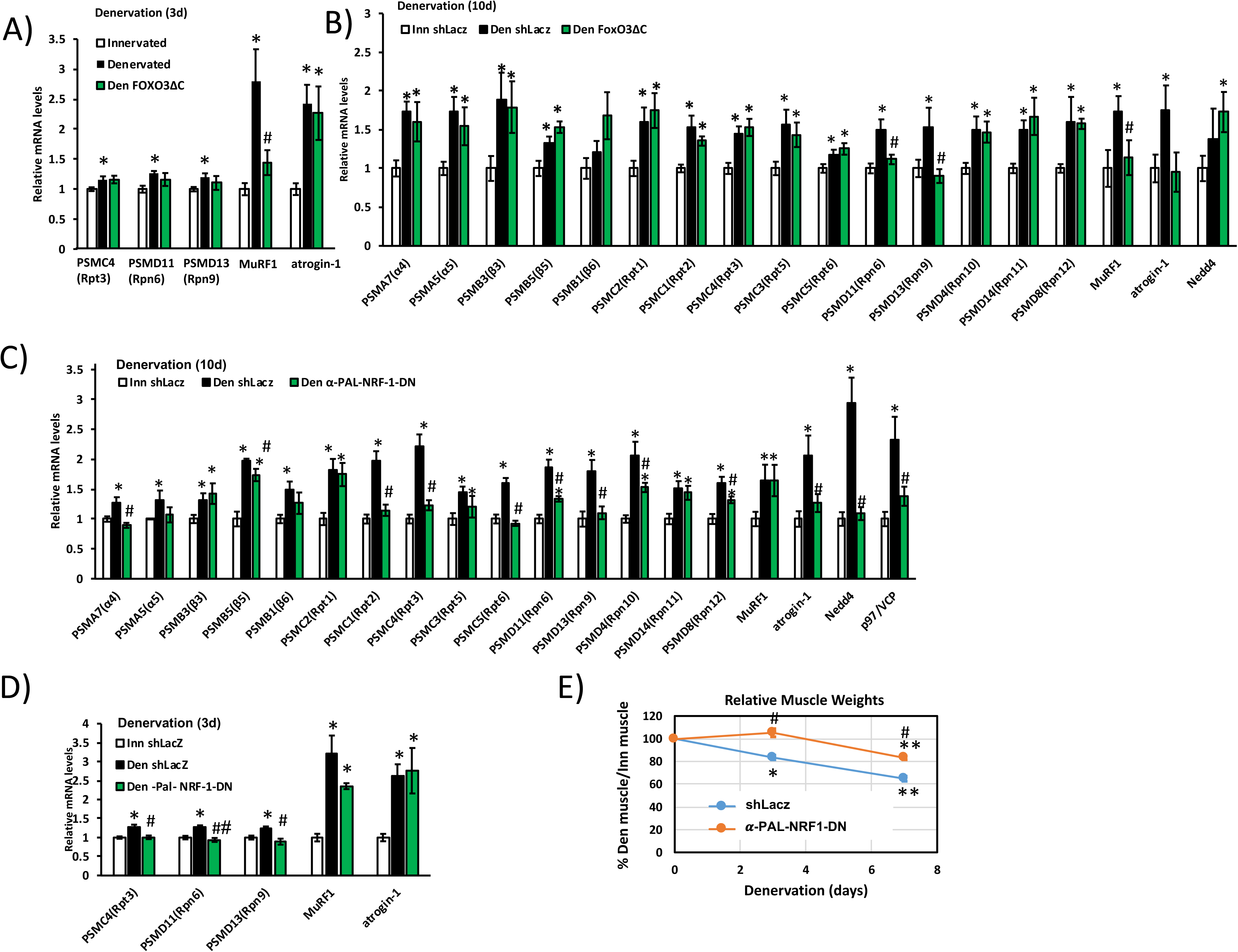
α-PAL^NRF-1^ is required for proteasome gene induction *in vivo*. A-B) Proteasome gene expression was measured by RT-PCR analysis of mRNA preparations from innervated and 3 (A) or 10 (B) days denervated muscles expressing dominant negative form of FOXO3 (FOXO3ΔC) or shLacz. FOXO3ΔC expression blocked the induction of two proteasome subunit genes (Rpn6 and Rpn9) at 10 d, but had no effect on proteasome gene induction at 3 d. N=4. *, P < 0.05 *vs.* innervated shLacz; # P < 0.05 *vs.* denervated shLacz. Error bars represent SEM. C-D) TA muscles were electroporated with a plasmid encoding α-PAL^NRF-1^-dominant negative (α-PAL^NRF-1^-DN) or shLacz and mRNA preparations from transfected muscles were analyzed by RT-PCR and specific primers at 10 (C) or 3 (D) days after denervation. N=5. * P < 0.05 *vs.* innervated; # P < 0.05 *vs.* denervated shLacz; ## P < 0.001 *vs.* denervated shLacz. Error bars represent SEM. E) Inhibition of NRF1 by the overexpression of α-PAL^NRF-1^-DN reduces atrophy of denervated muscles. Mean weights of denervated TA muscles are depicted as percent of innervated controls. N= 5 mice. * P < 0.05 and ** P < 0.001 *vs.* innervated; # P < 0.05 *vs.* denervated shLacz. Error bars represent SEM.

The transcription factor NRF-1^NFE2L1^ is a known regulator of proteasome gene induction in cultured cells (Radhakrishnan et al 2010b) but has not previously been studied in a whole organism *in vivo* or in atrophy. To test whether changes in NRF-1^NFE2L1^ levels can influence proteasome gene induction *in vivo*, we downregulated NRF-1^NFE2L1^ in TA muscle using a specific shRNA, and measured expression of proteasome genes and the atrogenes MuRF1 and atrogin-1 at 10 d after denervation. RT-PCR analysis on mRNA preparations from transfected muscles indicated that downregulating NRF-1^NFE2L1^ blocked the induction of certain proteasome genes (Fig. S3A), but not of MuRF1 and atrogin-1 (Fig. S3B).

Because ChIP sequencing data from cultured SK-N-SH human neuroblastoma cells have identified proteasome subunits as likely target genes of α-PAL^NRF-1^ ^24^, we sought to investigate the potential role of α-PAL^NRF-1^ in inducing proteasome genes *in vivo*. We searched the promoter regions of proteasome subunits for potential α-PAL^NRF-1^ (gene name: *NRF1*) binding sites using Fimo algorithm from the MEME suit ^38^ (Table 2). Potential α-PAL^NRF-1^ binding sites were chosen based on significant similarity (p< 0.05) to the human α-PAL^NRF-1^ motif (**<u>MA0506.1</u>)** ^38^. Among all proteasome genes, 13 contained potential binding sites for α-PAL^NRF-1^ (Table 2). Interestingly, 12 of these 13 genes contained binding sites for both α-PAL^NRF-1^ and PAX4 (Table 2).

To determine the role of α-PAL^NRF-1^ in atrophying muscle, we suppressed its function in denervated muscle by the electroporation of a plasmid encoding an α-PAL^NRF-1^ dominant negative (α-PAL^NRF-1^-DN) into TA muscles. The protein α-PAL^NRF-1^ binds DNA as a homodimer, and the dominant negative lacks the transactivation domain, thereby preventing dimerization and inhibiting the endogenous enzyme ^44, 45^. Proteasome genes were induced at 10 d after denervation in muscles expressing shLacz control (Fig. 5C). However, expression of the α-PAL^NRF-1^-DN blocked the induction of 9 proteasome subunits (out of 15 genes that were tested), two proteasome chaperones, and the atrophy-related genes atrogin-1, Nedd4, and p97/VCP (Fig. 5C).

The transcription factor α-PAL^NRF-1^ seems to be important to sustain proteasome gene expression throughout the process of atrophy because its inhibition with α-PAL^NRF-1^-DN also at the early phase (i.e. at 3 d after denervation) blocked the induction of PSMC4 (Rpt3), PSMD11 (Rpn6), and PSMD13 (Rpn9) (Fig. 5D). By contrast, whether or not α-PAL^NRF-1^ expression was inhibited, the ubiquitin ligases MuRF1 and atrogin-1 were induced in 3 d denervated muscle compared to innervated controls (Fig. 5D), probably because their expression in the early phase is primarily controlled by FOXO ^20^. Thus, α-PAL^NRF-1^ is essential for proteasome gene induction in denervated muscle, which promotes most of the accelerated proteolysis in atrophying muscles. In fact, denervation caused a gradual decrease over time in mean weights of TA muscles, but overexpression of α-PAL^NRF-1^-DN markedly attenuated this wasting (Fig. 5E), indicating that α-PAL^NRF-1^ function is important for proteolysis and loss of muscle mass. Thus, α-PAL^NRF-1^ is a novel transcription factor required for proteasome gene expression and overall proteolysis *in vivo*.

### Proteasome gene expression is dependent on both PAX4 and α-PAL^NRF-1^

These studies demonstrated that on denervation, induction of several genes that promote proteolysis is dependent on more than one transcription factor. Downregulation of either NRF-1 or PAX4 blocked PSMC1(Rpt2) expression (Figs. 4 and 5), downregulation of either NRF-1 or FOXO3 blocked PSMD11 (Rpn6) and PSMD13 (Rpn9) expression (Fig. 5), and induction of the bona fide atrophy markers, both MuRF1 and atrogin-1, required more than one transcription factor: MuRF1 expression was controlled by PAX4 and FOXO3, and atrogin-1 by PAX4, FOXO3, and α-PAL^NRF-1^. Thus, gene induction of proteasome subunits and UPS components in catabolic states *in vivo* appears to occur via the cooperative functions of multiple transcription factors.

To further study the potential cooperativity between PAX4 and α-PAL^NRF-1^, we generated an inducible α-PAL^NRF-1^/PAX4 double whole-body knockdown (KD) mouse (Fig. 6A). Initially, we generated a α-PAL^NRF-1^ whole body KO mouse, which had lower body weight and smaller muscles, and died (40% of progeny) a few weeks after birth (Fig. 6B and C). Thus, α-PAL^NRF-1^ is likely serving a crucial function in vital organs. We therefore generated a heterozygous Cre^+^/α-PAL^NRF-1^(fl/+) KD mouse, which was then crossed with PAX4 KO mice to generate α-PAL^NRF-1^/PAX4 double KD mouse (Fig. 6A). Cre^+^/α-PAL^NRF-1^(fl/+) KD and α-PAL^NRF-1^/PAX4 double KD mice showed normal body weights and survival (Fig. 6B and C) and no gross abnormalities compared to WT littermates.

**Figure 6.**
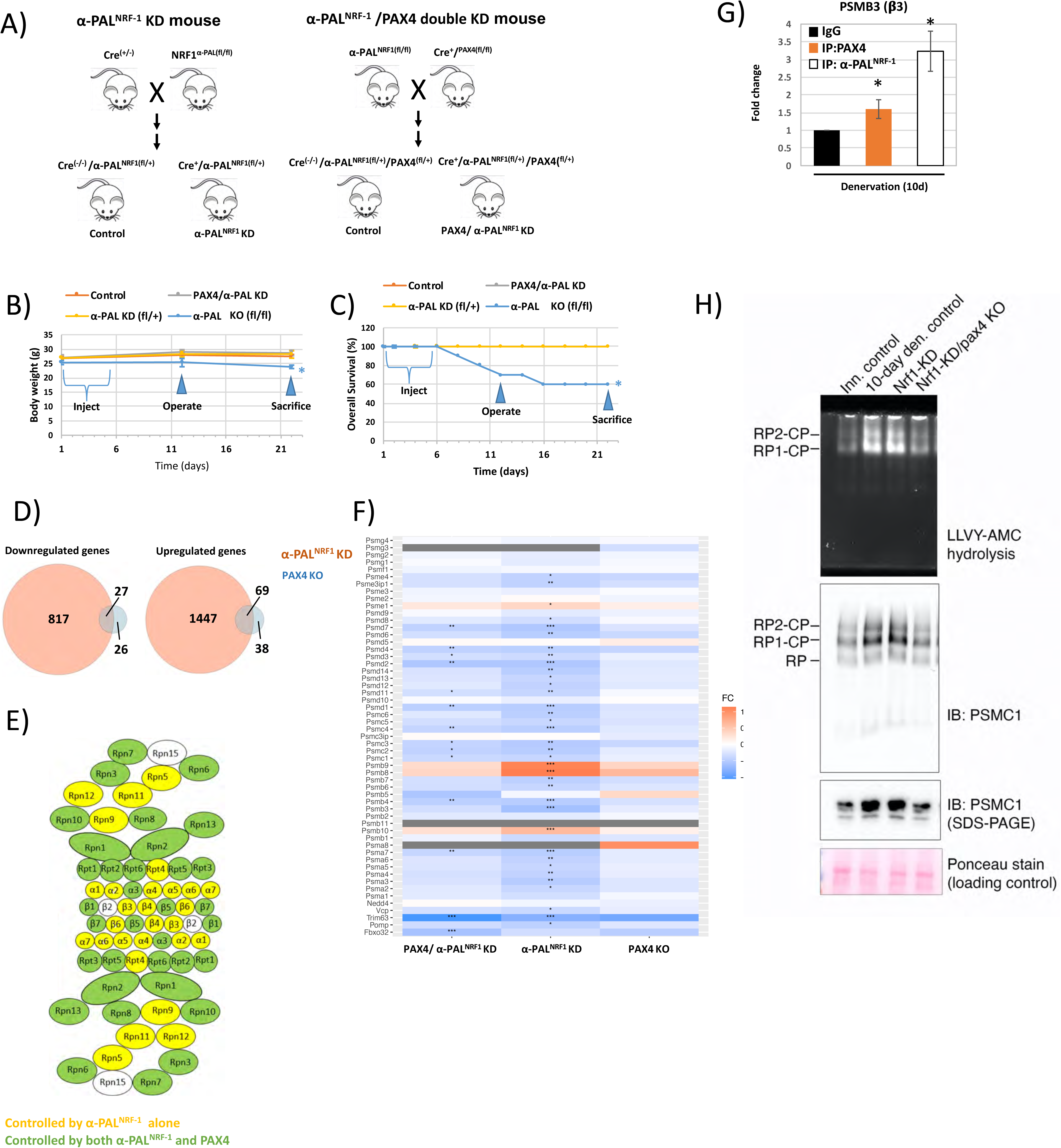
Proteasome gene expression is dependent on both PAX4 and α-PAL*^NRF-1^*. A) Strategy for generating α-PAL^NRF-1^ and PAX4/ α-PAL^NRF-1^ double conditional KD mice. Following tamoxifen injections, removal of α-PAL^NRF-1^ from one allele in the genome was verified using the U3/U5 primers (Supplemental Table S3). B-C) Body weight (g)(B) and survival (%)(C) of PAX4 KO, α-PAL^NRF-1^ KD, PAX4/α-PAL^NRF-1^ double conditional KD, and WT mice are presented, following tamoxifen injections and muscle denervation. D) Venn dagrams dipicting the overlap between differentially expressed genes (DEGs) upon α-PAL^NRF-1^ KD and DEGs upon PAX4 KO following tamoxifen injections and muscle denervation. A significant overlap was detected for the downregulated genes sh (p=2.5e-29, Fisher Exact test) and the upregulated genes (p=3.3e-266, Fisher Exact test) . E) Denervated muscles (10 d) were compared to innervated muscles by RNA sequencing. Presented are proteasome subunits whose induction was blocked in α-PAL^NRF-1^ KD (yellow) and PAX4/α-PAL^NRF-1^ double conditional KD (green) mice, with the latter group of genes being fully contained within the first. F) A heatmap representing the calculated Fold Changes (FCs) of the expression of the proteosome genes upon PAX4 and α-PAL^NRF-1^ double KD (left column), α-PAL^NRF-^1 (middle column) and PAX4 KO (Right column). Asterix denote significant FCs (* padj range 0.05-0.01, ** padj 0.01-0.001, *** padj < 0.001) G) PAX4 and α-PAL^NRF-1^ bind the promoter region of *PSMB3* gene. ChIP was performed on denervated (10 d) muscles from WT mice using PAX4 or α-PAL^NRF-1^ antibodies, or non-specific IgG control, and primers for *PSMB3* gene. Data is plotted as fold change relative to IgG control. N=3. *, P < 0.05 *vs.* IgG control. Error bars represent SEM. H) The content of active assembeled proteasomes increase in denervated muscles, but not in msucles lacking both PAX4 and α-PAL^NRF-1^. Measurement of proteasome content by native gels and immunoblotting, and proteasome peptidase activity by LLVY-AMC cleavage.

To investigate the coordinated functions of PAX4 and α-PAL^NRF-1^ in controlling proteasome gene expression, we performed RNA-seq of mRNA preparations from 10 day denervated muscles, when most proteasome genes and related chaperones are induced (Fig. 2), proteasome assembly is high (Fig. 3A), and myofibrils disassemble ^19, 26, 34^. We initally compared samples from innervated muscles, 10 d denervated muscles, and 10d denervated PAX4 KO muscles. For rigorous statistical analysis, we used p adjusted values that take into account the the false discovery rate to evaluate significance (p_adj_ < 0.05). Of the 38 proteasome genes detected by RNA-Seq, 22 were upregulated in denervated muscles *vs.* innervated controls (Table 3). The upregulated genes encoded multiple α and β 20S subunits, Rpn subunits, and Rpt subunits, in agreement with our findings above. Among these genes were PSMC1-6 (all Rpt subunits), PSMD4/8/11/13/14, and PSMA5/7, and PSMB1/3/5 (Fig. 2). By cotrast, PSME1 (PA28α) was downregulated, in agreement with our data shown in Fig. 2I. The remaining 16 proteasome subunit genes showed no statistically significant change (p_adj_ > 0.05), but two of them (PSMA2 and PSMD7) showed a similar trend toward higher levels in denervated muscles (p-value < 0.05). Other UPS and atrophy-related genes were also detected by RNA-seq, including proteasome activity regulators (PA28β, PA28ψ, PA28G, PA200, PI31), proteasome assembly chaperons (PAC1, PAC2, PAC3, PAC4, POMP), MuRF1 and atrogin-1. As expected, MuRF1 and POMP were induced (p_adj_ < 0.05), while atrogin-1 showed a similar trend (p < 0.05, but p_adj_ > 0.05). PAX4 deficiency in muscles from PAX4 KO mice showed a trend of reduction (p < 0.05, though p_adj_ > 0.05) in the expression of several proteasome genes including PSMD4 (Rpn10), PSMD6 (Rpn7), PSMD12 (Rpn5), PSMC3 (Rpt5), PSMC5 (Rpt6), PSMA3 (α7), PSMB7 (ý7), PSMA2 (α2), and of MuRF1 and atrogin1 (Table 3).

**Table 3.** RNA-seq data statistics for proteasome and UPS genes in WT innervated controls *vs*. WT or PAX4 KO denervated muscles (10 d).

|  | Denervated vs. Innervated |  |  | PAX4 KO denervated vs. WT |  |
| --- | --- | --- | --- | --- | --- |
| Gene | log2FoldChange | p-value | padj | log2FoldChange | p-value |
| Psma1 | 0.376139399 | 0.000793058 | 0.004894715 | -0.081824601 | 0.462402749 |
| Psma2 | 0.294110622 | 0.020357345 | 0.070522615 | -0.248274341 | 0.049258297 |
| Psma3 | 0.370898928 | 0.004291509 | 0.020023883 | -0.247794637 | 0.055164788 |
| Psma4 | 0.62502165 | 3.68E-05 | 0.00035577 | -0.17041028 | 0.25824041 |
| Psma5 | 0.529339637 | 0.00018006 | 0.001409121 | -0.155805072 | 0.264629117 |
| Psma6 | 0.216348708 | 0.092908763 | 0.223377373 | -0.192361093 | 0.134104149 |
| Psma7 | 0.749034613 | 8.45E-06 | 0.000100102 | -0.192819606 | 0.249630995 |
| Psma8 | 0.685340067 | 0.55945904 | 0.734881638 | 1.006686777 | 0.31413066 |
| Psmb1 | 0.767966033 | 3.40E-07 | 5.71E-06 | -0.244073074 | 0.103638508 |
| Psmb10 | 0.073184615 | 0.740874184 | 0.858606896 | 0.127058635 | 0.562130989 |
| Psmb2 | 0.041468297 | 0.703163425 | 0.835139258 | -0.077081451 | 0.476574817 |
| Psmb3 | 0.679355361 | 0.000248078 | 0.001834885 | -0.179996582 | 0.329138084 |
| Psmb4 | 0.647766184 | 0.000150997 | 0.001207162 | -0.285175986 | 0.094266441 |
| Psmb5 | 0.609916658 | 0.065839103 | 0.172824283 | 0.344036308 | 0.296517052 |
| Psmb6 | 0.407895698 | 0.007135007 | 0.030399481 | -0.244751078 | 0.105550925 |
| Psmb7 | 0.200873618 | 0.197519486 | 0.378975523 | -0.326702356 | 0.035971952 |
| Psmb8 | -0.145502571 | 0.46020198 | 0.655028294 | 0.634940494 | 0.000915478 |
| Psmb9 | 0.137195436 | 0.630457815 | 0.787000479 | 0.36994247 | 0.182372515 |
| Psmc1 | 0.759723724 | 3.88E-07 | 6.44E-06 | -0.150393019 | 0.312562672 |
| Psmc2 | 0.527783558 | 1.53E-05 | 0.000168728 | -0.18907803 | 0.119612365 |
| Psmc3 | 0.714272306 | 1.25E-05 | 0.000140738 | -0.386784881 | 0.017795087 |
| Psmc4 | 0.55269472 | 0.000193792 | 0.00149354 | -0.21630929 | 0.142733116 |
| Psmc5 | 0.460493029 | 0.000315613 | 0.002250713 | -0.270302958 | 0.033645275 |
| Psmc6 | 0.620835182 | 1.39E-06 | 2.00E-05 | -0.246302638 | 0.054430551 |
| Psm d1 | 0.199495859 | 0.160437845 | 0.328249599 | -0.265020109 | 0.061836411 |
| Psm d10 | 0.168386841 | 0.285080785 | 0.48443376 | 0.006802275 | 0.964652615 |
| Psm d11 | 0.951327366 | 5.78E-15 | 4.22E-13 | -0.066434381 | 0.581611501 |
| Psm d12 | 0.184159705 | 0.086315297 | 0.211589388 | -0.215638019 | 0.043891239 |
| Psm d13 | 0.535971439 | 1.91E-05 | 0.000202492 | -0.18956662 | 0.127858082 |
| Psm d14 | 0.490268234 | 6.65E-05 | 0.000595231 | -0.202424133 | 0.096432904 |
| Psm d2 | 0.425759359 | 0.004375947 | 0.020309747 | -0.225708386 | 0.130258794 |
| Psm d3 | 0.301899911 | 0.061659181 | 0.164811921 | -0.139096454 | 0.388052958 |
| Psm d4 | 1.05123923 | 7.17E-10 | 2.21E-08 | -0.405767467 | 0.016988007 |
| Psm d5 | 0.532146052 | 1.72E-05 | 0.000186443 | 0.155812992 | 0.191594248 |
| Psm d6 | 0.370134735 | 0.005423814 | 0.024278684 | -0.37711163 | 0.004515567 |
| Psm d7 | 0.332107946 | 0.019199542 | 0.067395954 | -0.263372101 | 0.062755971 |
| Psm d8 | 0.909206011 | 1.24E-05 | 0.00014023 | -0.053198799 | 0.797744136 |
| Psm d9 | 0.002912127 | 0.985694378 | 0.991986125 | -0.124318489 | 0.441205556 |
| Psme1 (PA28 ) | -0.368922299 | 0.001954738 | 0.010518588 | 0.141254683 | 0.234504295 |
| Psme2 (PA28 ) | 0.493039747 | 0.003006596 | 0.015032982 | -0.104832779 | 0.522563368 |
| Psme3 (PA28 ) | 0.017875419 | 0.870571737 | 0.930221755 | -0.02636632 | 0.808604085 |
| **Psme3ip1 (PA28G)** | 0.494063814 | 0.001276154 | 0.007347036 | -0.301506569 | 0.046716667 |
| **Psme4 (PA200)** | -0.291519722 | 0.134302244 | 0.289885303 | -0.379895988 | 0.051154884 |
| **Psmf1 (PI31)** | -0.055377576 | 0.733868688 | 0.85415032 | -0.033988596 | 0.833441461 |
| **Psmg1 (PAC1 assembly chaperon)** | 0.275038141 | 0.1497535 | 0.312461231 | -0.066537212 | 0.721866993 |
| **Psmg2 (PAC2 assembly chaperon)** | 0.481145132 | 0.000945686 | 0.005671471 | -0.096660452 | 0.496666029 |
| **Psmg3 (PAC3 assembly chaperon)** | 0.544537182 | 0.664850092 | 0.810161116 | -0.319644871 | 0.790314505 |
| **Psmg4 (PAC4 assembly chaperon)** | 1.023431913 | 6.30E-05 | 0.000569276 | -0.054210618 | 0.822945429 |
| **Trim63** | 0.76423456 | 0.007979885 | 0.033271099 | -0.88452467 | 0.002134156 |
| **Fbxo32** | 0.542463241 | 0.014169018 | 0.052857372 | -0.480586907 | 0.029725298 |
| **Pomp** | 0.768928645 | 3.24E-07 | 5.45E-06 | -0.239400181 | 0.109321305 |

| denervated |
| --- |
| padj |
| 0.822318892 |
| 0.391729305 |
| 0.409454263 |
| 0.693565291 |
| 0.700904206 |
| 0.558532439 |
| 0.686445749 |
| NA |
| 0.513021954 |
| 0.87081631 |
| 0.827876333 |
| 0.745842182 |
| 0.498214144 |
| 0.727182925 |
| 0.51578311 |
| 0.350370486 |
| 0.069471605 |
| 0.623339892 |
| 0.736476292 |
| 0.537282319 |
| 0.265325966 |
| 0.572020025 |
| 0.342127585 |
| 0.407774657 |
| 0.426997057 |
| 0.990231263 |
| 0.877895457 |
| 0.37699057 |
| 0.548756951 |
| 0.501348018 |
| 0.552052622 |
| 0.785113645 |
| 0.260420171 |
| 0.63377526 |
| 0.144284627 |
| 0.43049428 |
| 0.951888547 |
| 0.813156945 |
| 0.672340021 |
| 0.849297715 |
| 0.954417598 |
| 0.385399273 |
| 0.395937198 |
| 0.960104959 |
| 0.930202406 |
| 0.839039214 |
| NA |
| 0.9579293 |
| 0.10140899 |
| 0.324785076 |
| 0.518017514 |

A genome wide comparison using RNA-seq of differentiatlly expressed genes in muscles from PAX4 KO and α-PAL^NRF-1^ KD mice indicated that α-PAL^NRF-1^ controls the expression of hundreds of genes at 10 d after denervation, while the effects of PAX4 are on a few dozens of genes (Fig. 6D). Interestingly, 27 genes are downregulated and 69 genes are upregulated by both PAX4 and α-PAL^NRF-1^ (Fig. 6D). In addition, the 27 genes that are induced by both transcription factors represent 50% of the genes controlled by PAX4, and the 69 genes that are downregulated by both transcription factors represent 64% of the genes controlled by PAX4 (Fig. 6D). These data, together with the presence of binding sites for both α-PAL^NRF1^ and PAX4 in 12 of the 13 proteasome genes, suggested that these two transcription factors may act cooperatively, possibly coregulating proteasome gene expression.

Therefore, we compared by RNA-seq samples from 10d denervated muscles to 10d denervated α-PAL^NRF-1^ KD or PAX4/ α-PAL^NRF-1^ double KD muscles. We found that the expression of 30 proteasome genes decreased in α-PAL^NRF-1^ KD muscles compared with denervated muscles from WT littermates (Table 4), and PSMB1 showed a similar trend (p < 0.05, but p_adj_ > 0.05). In PAX4/ α-PAL^NRF-1^ double KD mouse muscles the expression of 12 proteasome genes decreased (p_adj_ < 0.05), and additional 12 genes showed a similar trend of reduction (p < 0.05, but p_adj_ > 0.05). As expected, the group of genes affected by PAX4/ α-PAL^NRF-1^ double KD was fully contained within the group of genes affected by α-PAL^NRF-1^ KD alone (Table 4 and Fig. 6E and F). Overall, PAX4 KD seems to have a moderate effect on α-PAL^NRF-1^ KD regulation of proteasome genes (Table 4 and Fig. 6F).

**Table 4.**
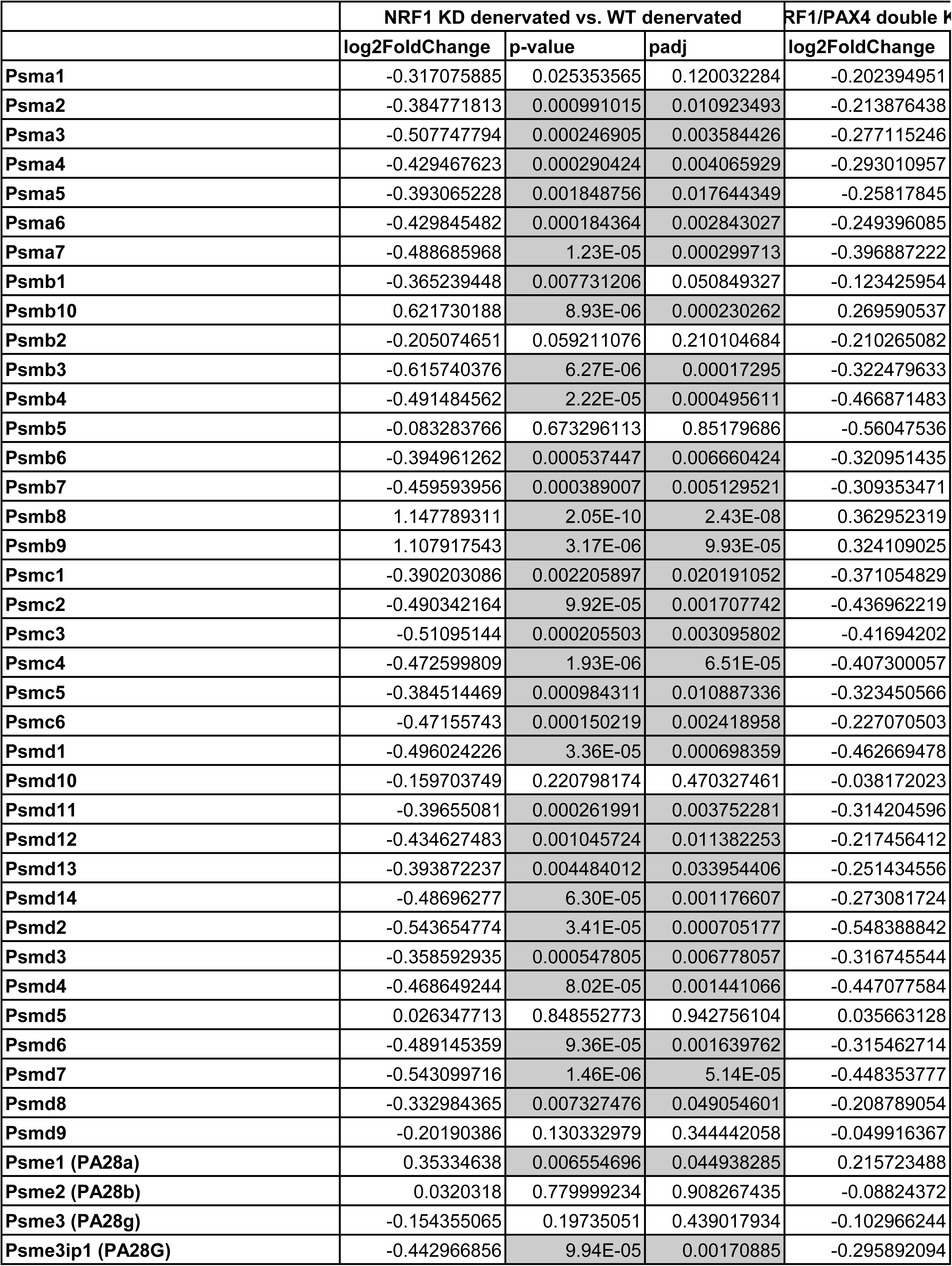
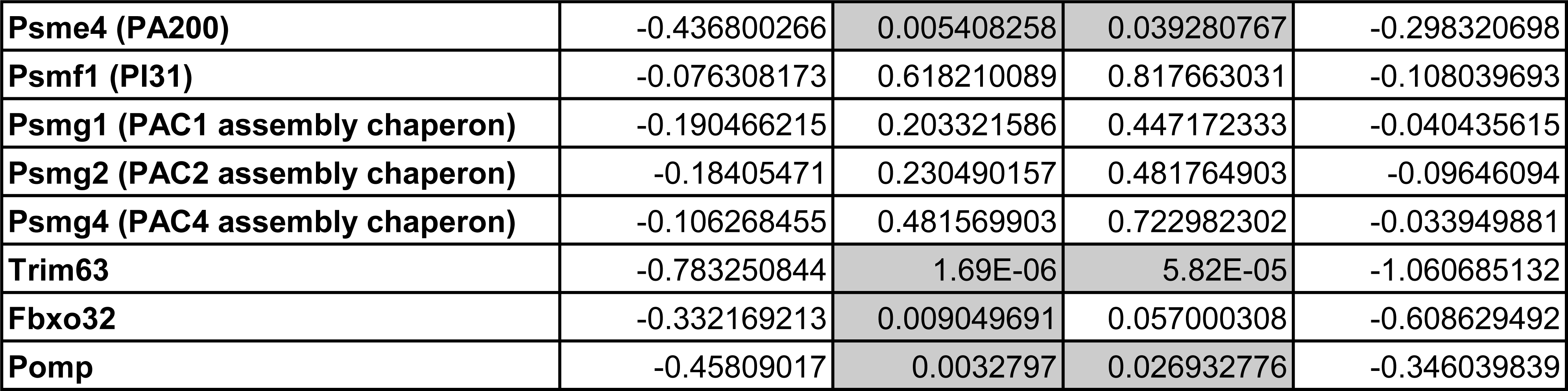
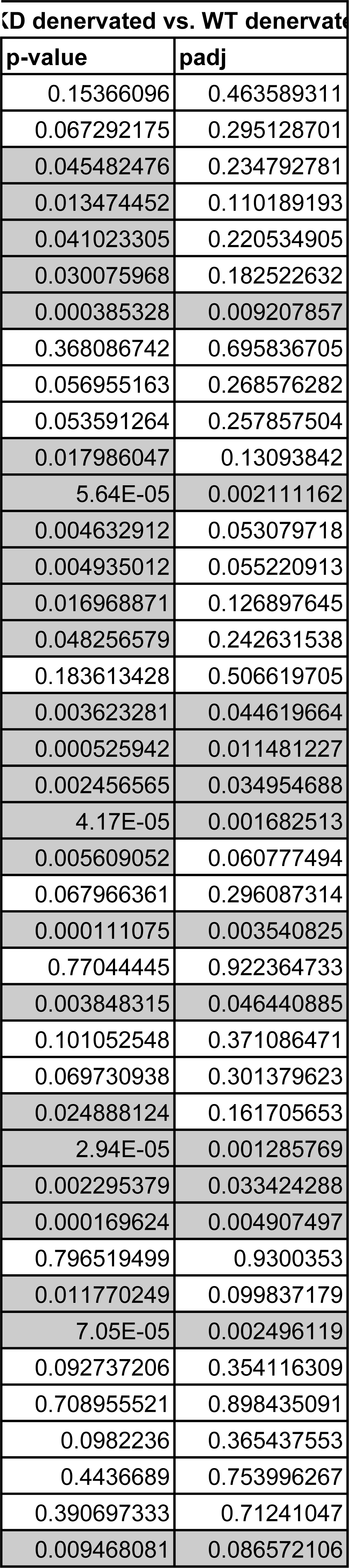
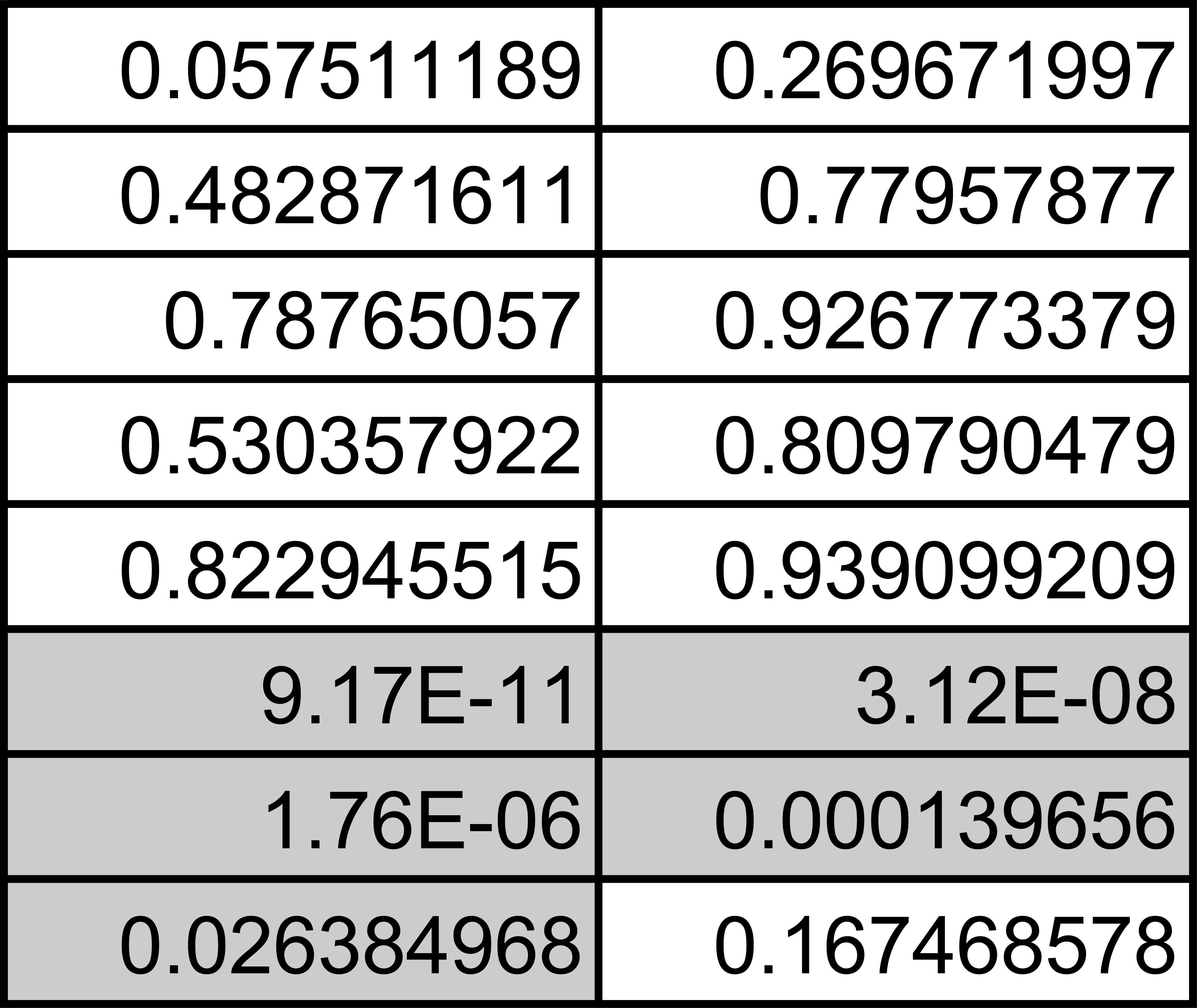
RNA-seq data statistics for proteasome and UPS genes in WT denervated (10 d) *vs.* α-PAL^NRF-1^ KD or PAX4/ α-PAL^NRF-1^ double KD muscles.

Together, our analyses showed a significant overlap between genes that were significantly downregulated and those that were upregulated upon PAX4 KO, α-PAL^NRF-1^ KD or PAX4/ α-PAL^NRF-1^ double KD (Fisher Exact test; p=2.3e-29 and p=3.3e-66 for downregulated and upregulated, respectively) (Fig 6D). These findings suggest the two transcription factors may act together to regulate the same set of genes, which was validated for a representative gene, PSMB (ý3), by ChIP (Fig. 6G). Moreover, analysis of innervated and 10 d denervated muscle homogenates from WT, α-PAL^NRF-1^ KD or PAX4/ α-PAL^NRF-1^ mice by native gels and immunoblotting or LLVY-cleavage indicated that loss of both transcription factors is necessary to effectively block accumulation of active assembled proteasomes on denervation (Fig. 6H). Although knock-down of α-PAL^NRF-1^ was sufficient to reduce proteasome gene expression (Fig. 6F), it did not block the increase in active proteasomes on denervation (Fig. 6H), probably because PAX4 was still present in those samples, and the presence of both transcription factors is necessary to boost proteasome content in atrophying muscle.

## DISCUSSION

Here we discovered a novel mechanism that regulates proteasome gene expression in mouse muscle *in vivo*, involving an unprecedented functional dependency between two transcription factors, PAX4 and α-PAL^NRF-1^. The structure, regulation and assembly of the proteasome are well studied *in vitro* in yeast and in mammalian cell cultures (Bard et al., 2018; Budenholzer et al., 2017), but significantly less so *in vivo* in mammals ^46, 47^. Thus, our studies uncover for the first time the mode of regulation of proteasome gene expression *in vivo*, using denervated adult mouse muscle as a model system.

Muscle atrophy is a valuable physiological system to study protein degradation by the proteasome in a whole organism *in vivo*. It had long been known that this catabolic condition is associated with severe maladies such as denervation, cancer, chronic inflammation, aging, neurodegeneration, and malnutrition ^12^. The inevitable loss of muscle mass and strength results from protein breakdown outstripping synthesis, leading to frailty, disability, morbidity and mortality. During times of scarcity or illness, this survival process is designed to guarantee the continued availability of amino acids for energy production by the liver, to nurture vital tissues such as the brain and heart ^46^. Proteolysis upon muscle denervation appears to take place at two stages, which may be readily observed due to its relatively slow pace compared to starvation or inflammation driven atrophy ^19^. During the first stage of atrophy, the major atrogenes MuRF1 and atrogin1 are induced, and in the latter stage there is an increase in other genes that promote proteolysis (e.g. NEDD4, RPT1, p97/VCP) to support the accelerated degradation of myosin and actin and associated myofibrillar proteins ^19, 26, 35^. Our findings here support this concept of coordinated early and late phases of gene induction during atrophy: in the early phase (3 – 7 d after denervation), genes encoding 20S subunits, POMP, atrogin-1, MuRF1, and Rpn subunits are induced. In a later phase (10 – 14 d after denervation), Nedd4, p97/VCP, and Rpn subunits (again) are induced. These findings suggest that basal proteasome levels are sufficient to carry out the muscle loss that occurs early after denervation (3 - 7 d after denervation). Then, proteasome subunits and chaperones are elevated at 7 d, prior to the rapid myofibril disassembly that occurs in the late phase ^26^. Late in atrophy (14 d after denervation), levels of most genes returned toward basal levels, as shown by RT-PCR and RNA-seq data. This is coordinated with a slowing in the rate of loss of muscle and myofibril content ^11^.

Many of the major atrogenes are induced by FOXO family of transcription factors ^48^; their roles are well established during the early stages of atrophy (1 d fasting or 3 d denervation), and the induction of FOXO3 alone is sufficient to induce skeletal muscle atrophy without further stimuli ^20^. The role of FOXO3 in the later stages of atrophy (10d denervation), where further induction of many proteasome genes is observed concomitantly with accelerated proteolysis, was not well documented in the literature, and here we show it induces PSMD11 (Rpn6), PSMD13 (Rpn9), and MuRF1 (Fig. 5B). In addition, a recent study from our lab demonstrated that PAX4 is a novel transcription factor for the proteasome gene PSMC2 (Rpt1) during the later stages of atrophy ^19^. As most proteasome components are induced in skeletal muscle cells at this stage, it is expected that other transcription factors must play a role too.

NRF-1^NFE2L1^ is one of the few transcription factors that has been shown in cultured cells to regulate proteasome genes, through binding to CCAAT boxes in gene promoters^49^. Its most notable targets are STAT3, and several β proteasome subunits ^50^, while STAT3 has been linked to muscle wasting caused by denervation ^51^ and cancer cachexia^50^. Our studies in mouse muscle now reveal what transcription factors in fact regulate proteasome gene expression *in vivo*. In line with prior investigations in cultured MEF cells^8^, we show here that downregulation of NRF-1^NFE2L1^ in mouse muscle *in vivo* blocks the induction of the proteasome genes PSMA7 (α4), PSMB4 (ý7), PSMC1 (Rpt2), PSMC4 (Rpt3)(Fig. S3A). We provide evidence for additional proteasome genes that are controlled by this transcription factor *in vivo*, including PSMC3 (Rpt5) and PSMC5 (Rpt6). However, our data indicate an even stronger suppressive effect on most proteasome genes in denervated muscles by the inhibition (using dominant negative) or knock-down in transgenic mice of α-PAL^NRF-1^, which was also sufficient to attenuate muscle atrophy (Fig. 5E). Such role for α-PAL^NRF-1^ in inducing proteasome genes has not been documented before. Furthermore, we identified potential binding sites for this transcription factor in many proteasome genes, which was validated for a representative gene by ChIP (Fig. 6G). Moreover, RNA-seq of denervated muscles from α-PAL^NRF-1^ knock-down mice showed a global effect on proteasome and UPS gene expression (Fig. 6E-F). Thus, these data indicate a physiological role for α-PAL^NRF-1^ in promoting muscle loss after denervation by inducing proteasome subunits and UPS components.

Surprisingly, the expression of several proteasome subunit genes is controlled by both PAX4 and α-PAL^NRF-1^, suggesting that proteasome gene induction in atrophy is regulated by multiple transcription factors. Likely, their roles are not redundant as loss of one transcription factor is sufficient to prevent induction of various proteasome genes in muscle after denervation. Our data on proteasome assembly provide further insights on how these two transcription factors might regulate proteasome subunit genes. Although both transcription factors can induce proteasome subunit genes, mainly PAX4 KO attenuated the increased assembly of proteasome (Fig. 6H). In addition, none of these transcription factors controls the expression of the proteasome assembly chaperon genes PSMD5, PSMD9, and PSMD10 (Fig. S4). These results suggest that PAX4-induced mechanism might facilitate a more stochiometric production of individual subunits, thereby synergistically increasing proteasome assembly, together with its assembly chaperones. α-PAL^NRF-1^ can increase proteasome gene expression, but might depend on PAX4 to maintain balanced expression of proteasome genes. Such a scenario may explain a seemingly conflicting result, in which proteasome gene expression is reduced in α-PAL^NRF-1^ knock-down, but the assembled proteasome complexes remain increased, due to PAX4 maintaining subunit stoichiometry for increased proteasome assembly.

Because common proteolytic pathways are activated in diverse types of atrophy, targeting key factors in of these common mechanisms should be beneficial in treating many diseases. It remains to be determined whether the new transcriptional regulation of proteasome gene expression by multiple transcription factors is important in other types of atrophy, including cancer, aging or diabetes. Depletion of PAX4 or α-PAL^NRF-1^ significantly reduces atrophy induced by denervation. Consequently, these transcription factors represent potential novel therapeutic targets to treat muscle wasting.

## ACKNOWLEDGEMENTS

This project was supported by grants from the Israel Science Foundation (grant no. 1068/19) to S. Cohen, and NIH (grant no. R01GM127688) to S. Park. Additional funds were received from the Russell Berrie Nanotechnology Institute, Technion to S. Cohen, and the Zuckerman STEM Leadership Program to J.E.G. We are extremely thankful to the LS&E Microscopy Center at Technion for their assistance with the confocal microscope.

## AUTHOR CONTRIBUTIONS

J.E.G. performed all experiments. D.K., N.T., performed experiments presented in Figures 4I-J and 6G. T.L., Y.M.G performed the bioinformatic analyses. T.G. performed the analyses of assembled proteasomes. J.E.G., S.P, Y.M.G, S.C designed experiments, analyzed data, and wrote the paper.

## STAR METHODS

### Animal experiments

All animal procedures were performed in accordance with the ethics guidelines of the Israel Council on Animal Experiments and the Technion Inspection Committee on the Constitution of the Animal Experimentation. Specialized staff provided animal care in the institutional specific pathogen-free animal facility. For experiments on wild-type (WT) mice, adult, Hsd:ICR male mice (26-30 g, Envigo) were used, and Tibialis Anterior (TA) or Gastrocnemius (GA) muscles were analyzed. Muscle denervation was performed by sectioning the sciatic nerve. At the same time, electroporation of TA muscles was performed as described ^37, 52^. Briefly, a plasmid for a specific shRNA (20 μg) (Supplemental Table S2) or α-PAL^NRF-1^ dominant negative (30 μg) in 0.9 w/v % NaCl was injected into the muscle, and five electric pulses (12V, 200 ms intervals) were applied. The α-PAL^NRF-1^ dominant negative plasmid was a kind gift from Dr. David Hockenberry at the Fred Hutchinson Cancer Research Center. Human HA-FOXO3ΔC (Addgene 1796) was used to inhibit FOXO3.

### Generation of transgenic mice

PAX4 KO, α-PAL^NRF-1^ knock-down (KD), and PAX4/α-PAL^NRF-1^ double KD mice were generated according to the breeding schemes in Figs. 4C and 6A. α-PAL^NRF-1fl/fl^ mice were kindly provided by Dr. Chai-An Mao, the University of Texas. PAX4^fl/+^ mice were generously provided by Dr. Ahmed Mansouri at the Max Planck Institute for Biophysical Chemistry ^41^. Cag-Cre mice were purchased from the Jackson Laboratory (Strain B6.Cg-Tg(CAG-cre/Esr1*)5Amc/J, Stock# 004682). DNA was extracted from tail snips taken at weaning or sacrifice for PCR genotyping (Taq DNA Polymerase, Ampliqon, Cat. #A180301). Genotyping primers are supplied in Supplemental Table S3. When male mice were 26-30 g (3-4 months), gene KO was induced by injecting 50 μL 20 mg/mL tamoxifen in corn oil per 10 g body weight every day for five days, followed by a one week wait period before conducting experiments.

### Muscle homogenization and fractionation

Muscles were homogenized in cold homogenization buffer (20 mM Tris pH 7.6, 5 mM EGTA, 100 mM KCl, 1% Triton X-100, 1 mM PMSF, 10 mM sodium pyrophosphate, 100 mM sodium fluoride, 2 mM sodium orthovanadate, 10 μg/mL aprotinin, 10 μg/mL leupeptin, 3 mM benzamidine hydrochloride) using an Omni Tissue Homogenizer, incubated with rotation at 4°C for 1 hr, and centrifuged at 6000 X g for 20 min at 4°C. The supernatant was removed as the soluble fraction. The pellet (insoluble fraction, Fig. 1C) was washed 1X with homogenization buffer and 2X with suspension buffer (20 mM Tris pH 7.2, 100 mM KCl, 1 mM DTT, 1 mM PMSF), then resuspended in storage buffer (20 mM Tris pH 7.2, 100 mM KCl, 1 mM DTT, 20% glycerol). Protein concentration was determined using Bradford Assay.

### Immunofluorescent staining and fiber size analysis

Frozen TA muscle cross-sections (30 μm) were fixed in 4% PFA in PBS, pH 7.4 for 20 min, washed and incubated overnight at 4°C in laminin antibody (Sigma, Cat. # L9393, 1:100), then washed and incubated in secondary antibody (anti-Rabbit IgG Alexa Fluor 568, Thermo Fisher Scientific, Cat. # A-11011, 1:500), followed by Hoechst staining for 5 min (Sigma, Cat. # H60241, 1:2500). For Wheat Germ Agglutinin (WGA Texas Red-X Conjugate, Invitrogen, Cat. # W21405, 1:100)), incubation was one hr at room temperature, with no secondary antibody. Confocal images were collected using an inverted LSM 710 laser scanning confocal microscope (Zeiss, Oberkochen, Germany) with a Plan-Apochromat 63 x 1.4 NA objective lens and BP 593/46, BP 525/50 and 640-797 filters. The fiber size area in muscle cross-sections was determined manually or using Imaris Image Analysis Software ^52^. For each comparison, equal numbers (>500) of muscle fibers were analyzed.

### Real-Time PCR

Total RNA was extracted from TA muscle using TRIzol reagent (Sigma, T9424), and served as a platform for cDNA synthesis using qPCRBIO cDNA Synthesis Kit (Cat. # PB30.11-10). Real-time PCR was performed with primers to mouse genes (Supplemental Table S2) using PerfeCTa SYBR Green FastMix ROX (Cat. # 95073) or qPCRBIO SyGreen Blue Mix Hi-ROX (Cat. # PB20.16-05) according to the manufacturer’s protocol. The gene Mrps7 was used as a reference. To amplify PAX4, which is very lowly expressed in muscle, RT-PCR was performed using one set of PAX4 primers (Table S2, set #1), and then 1µl of the product from this PCR reaction was amplified using a second set of PAX4 primers (Table S2, set #2) that is nested inside the first set. To reach the detection threshold for PAX4, 40 cycles were used.

### Chromatin Immunoprecipitation (ChIP) Assay

The ChIP assay was performed using the Millipore ChIP Assay Kit (Cat# 17-295). For each experimental condition, one Gastrocnemius muscle was lysed in Nuclear Extraction Buffer (10 mM HEPES pH 7.4, 10 mM KCl, 5 mM MgCl_2_, 0.5 mM DTT, protease and phosphatase inhibitors) using an Omni Tissue Homogenizer (for PAX4 immunoprecipitation from nuclear extracts, see protocol for tissue fractionation below). Samples were crosslinked with formaldehyde (1% v/v final concentration, 10 min, 4°C), quenched with 1.25 mM glycine for 5 min at room temperature, and centrifuged (1000 x g, 4°C, 5 min). Pellets were resuspended in SDS lysis buffer (Cat# 20-163) and subsequently sonicated with the Covaris E220 ultrasonicator (Peak Power 75, Duty Factor 26, Cycles/Burst 200, Temperature 6°C, 840 sec) to fragment DNA. The samples were then centrifuged at 1000 x g, 4°C for 5 min, and the supernatant was diluted with ChIP Dilution Buffer (Cat# 20-153), precleared for 1 hr at 4°C with 5 μl Protein A Agarose/Salmon Sperm DNA (Cat# 16-157C), and incubated with Protein A Agarose/Salmon Sperm DNA and 5 μg human IgG (Sigma, Cat# I4506), PAX4 Antibody (MyBioSource.com, Cat# MBS542431) or α-PAL^NRF-1^ antibody (DSHB Hybridoma Product PCRP-NRF-1-3H1. This antibody was deposited to the DSHB by Common Fund – Protein Capture Reagents Program) overnight at 4°C. The beads were washed with the buffers supplied in the kit in the following order for 7 min each: Low Salt Immune Complex Wash Buffer (Cat# 20-154), High Salt Immune Complex Wash Buffer (Cat# 20-155), LiCl Immune Complex Wash Buffer (Cat# 20-156), TE Buffer (Cat# 20-157). DNA was eluted in Elution Buffer (1% SDS, 0.1 M NaHCO_3,_ 0.2 mg/ml Proteinase K) for 5 hrs with shaking at 1200 rpm, 65°C. Eluted DNA was purified by isopropanol precipitation and subjected to RT-PCR analysis. Fold enrichment was calculated by dividing Cq values from the PAX4 or α-PAL^NRF-1^ immunoprecipitation by IgG signal.

Due to the low abundance of PAX4 in muscle, ChIP was performed on PAX4 precipitates from nuclear extracts that were prepared by tissue fractionation. Briefly, for each experimental condition one Gastrocnemius muscle was homogenized in 19 volumes (v/w) of buffer C (20 mM Tris, pH 7.6, 100 mM KCl, 5 mM EDTA, 1 mM DTT, 1 mM PMSF, 3 mM benzamidine, 10 μg/ml leupeptin, 50 mM NaF, and 1 mM sodium orthovanadate). Following centrifugation at 2,900 × g for 20 min at 4 °C, pellet was resuspended in 500ul Buffer C per 1mg of tissue, vortexed, and at 1000 g for 15 min at 4°C. Then, supernatant (cell debris) was discarded and the pellet was washed again in buffer C. The washed pellet was then resuspended in 200μl buffer N (20 mM HEPES pH 7.9, 1.5 mM MgCl_2_, 0.5 M NaCl, 5 mM EDTA/NAOH pH 7.4, 20% glycerol, 1% Triton X-100, 1 mM Sodium OrthoVanadate, 10 μg/ml Leupeptin, 3 mM Benzamidine, 1 mM PMSF, 50 mM NaF) per 50mg muscle, incubated on ice for 30 min, and passed 10–20 times through a 18-gauge needle. After centrifugation at 9,000 g for 30 min at 4°C, the supernatant (nuclear extract) was stored at -80°C until analysis.

### Computational Analysis

The promoter regions (1Kb upstream to the annotated Transcription Strat Sites) of the mouse proteasome genes and atrogenes were searched for potential transcription factor binding sites using the FIMO motif serach algorithm from the MEME suit (ref 43). The Position Specific Scoring Matrices (PSSMs) used to search the transcription factor binding site in the FIMO search tool were extracted from the JASPAR database of transcription factor binding profiles (ref PMID: 34850907). Genes in which a ignificant hit(p<0.05) was identified within its defined promoter region were considered as target genes.

### RNA sequencing

We trimmed low quality reads and removed ilumina adaptor and polyA sequences, using cutadapt ^53^. Reads were mapped against GRCm39 genome (Ensembl) using STAR package ^54^ to create Bam files. We used samtools to convert BAM to SAM file and index the files. Then, the reads were annotated and counted using htseq-count package^55^. Using Deseq package ^56^ we identified gene that are differentially expressed. R package was used to plot significant gene expression heatmaps and venn.

### Denaturing gel electrophoresis and Western blotting

Muscle homogenates were separated on 10 or 12.5% SDS polyacrylamide gels and transferred to PVDF membranes (2 hrs, 200 mA, 4°C). Membranes were blocked for 1 hr in 3% BSA/PBST, then incubated in primary antibodies overnight at 4°C, washed in PBST, and incubated in secondary antibody for 1 hr at room temperature. Membranes were then incubated 1 min in luminol ECL reagent and imaged using the Bio-Rad ChemiDoc. Antibodies and dilutions used were: PAX4: Abcam, Cat. # ab101721, 1:1000; Gankyrin (3A6C2): Santa Cruz, Cat. # sc-101498, 1:300; PSMD5 (S5b): Santa Cruz, Cat. # sc-390751, 1:100; Proteasome 20S α1, 2, 3, 5, 6 & 7 subunits (MCP231): Enzo, Cat. # BML-PW8195, 1:2500; PSMD13 (Rpn9): Proteintech, Cat. # 15261-1-AP, 1:2000; MuRF1: Regeneron, VCP (Clone 18/VCP): BD Biosciences; Laminin: Sigma, Cat. # L9393, 1:50; Rpt2: Bethyl laboratories, # A303-821A; Rpn1: Santa Cruz, # sc-271775; Peroxidase Goat Anti-Rabbit IgG (H+L): Jackson ImmunoResearch, Cat. # 111-035-144, 1:10000; Peroxidase Goat Anti-Mouse IgG (H+L): Jackson ImmunoResearch, Cat. # 115-035-003, 1:5000.

### Native-PAGE analysis of proteasome assembly and activity

Muscle tissues were ground with mortar and pestle in the presence of liquid nitrogen. The ground cryo-powders were hydrated for 10 min on ice using buffer A (50 mM Tris-HCl pH 7.5, 5 mM MgCl_2_, 75 mM NaCl, 0.5% NP-40, 10% glycerol) supplemented with 1 mM ATP and protease inhibitors. This buffer was added to the cryo-powders at a 1:1 ratio. The resulting cell lysates were centrifuged at 1500 x g for 20 min at 4°C to remove cell debris and insoluble proteins. The supernatant was centrifuged at 15,000 x g for 15 min twice at 4°C. The cleared lysates (100 mg) were loaded onto 3.5% discontinuous native gel. Native gels were electrophoresed for 3.5 hrs at 100V in the cold room. In-gel peptidase assays were conducted using the fluorogenic peptide substrate LLVY-AMC (Bachem, I-1395.0100) as described previously (we will insert reference here). Native gels were photographed under UV light Bio-Rad Gel Doc EZ Imager to detect AMC fluorescence. Following the in-gel peptidase assays, native-PAGE gels were transferred to PVDF membranes for analysis of proteasome assembly by immunoblotting. The PVDF membrane was incubated in 20 mL of blocking buffer, (TBST; Tris-buffered saline containing 0.1% Tween-20), which was supplemented with 5% non-fat dry milk, for 1 hr at room temperature. The membrane was washed twice for 10 min using TBST. The membrane was incubated with primary antibodies, which were diluted at 1:3000 in blocking buffer, overnight at 4°C. Two washes were conducted using TBST as above.

The HRP-conjugated secondary antibody was diluted at 1:3000 in blocking buffer (Cytiva, NA934, anti-Rabbit IgG HRP-linked antibody) for incubation with the membranes for 1 hr at room temperature, followed by two washes using TBST. Membranes were then subjected to Enhanced chemiluminescence (Perkin Elmer, Western Blot Chemiluminescence Reagents Plus) and were imaged using Bio-Rad ChemiDoc MP Imager.

### Statistical analysis and image acquisition

All data are presented as means ± SEM. The statistical significance was accessed with one-tailed unpaired Student’s *t* test. Muscle sections used for fiber size analysis were imaged at room temperature with an inverted LSM 710 laser scanning confocal microscope (Zeiss, Oberkochen, Germany) with a Plan-Apochromat 63 x 1.4 NA objective lens and BP 593/46, BP 525/50 and 640-797 filters. Image acquisition was performed using 3.2 ZEN imaging software (Zeiss), and data processing was performed using Imaris (Bitplane) software. Black and white images were processed with Adobe Photoshop 2021, version 22.4.2.

## SUPPLEMENTAL INFORMATION TITLES AND LEGENDS

**Figure S1.**
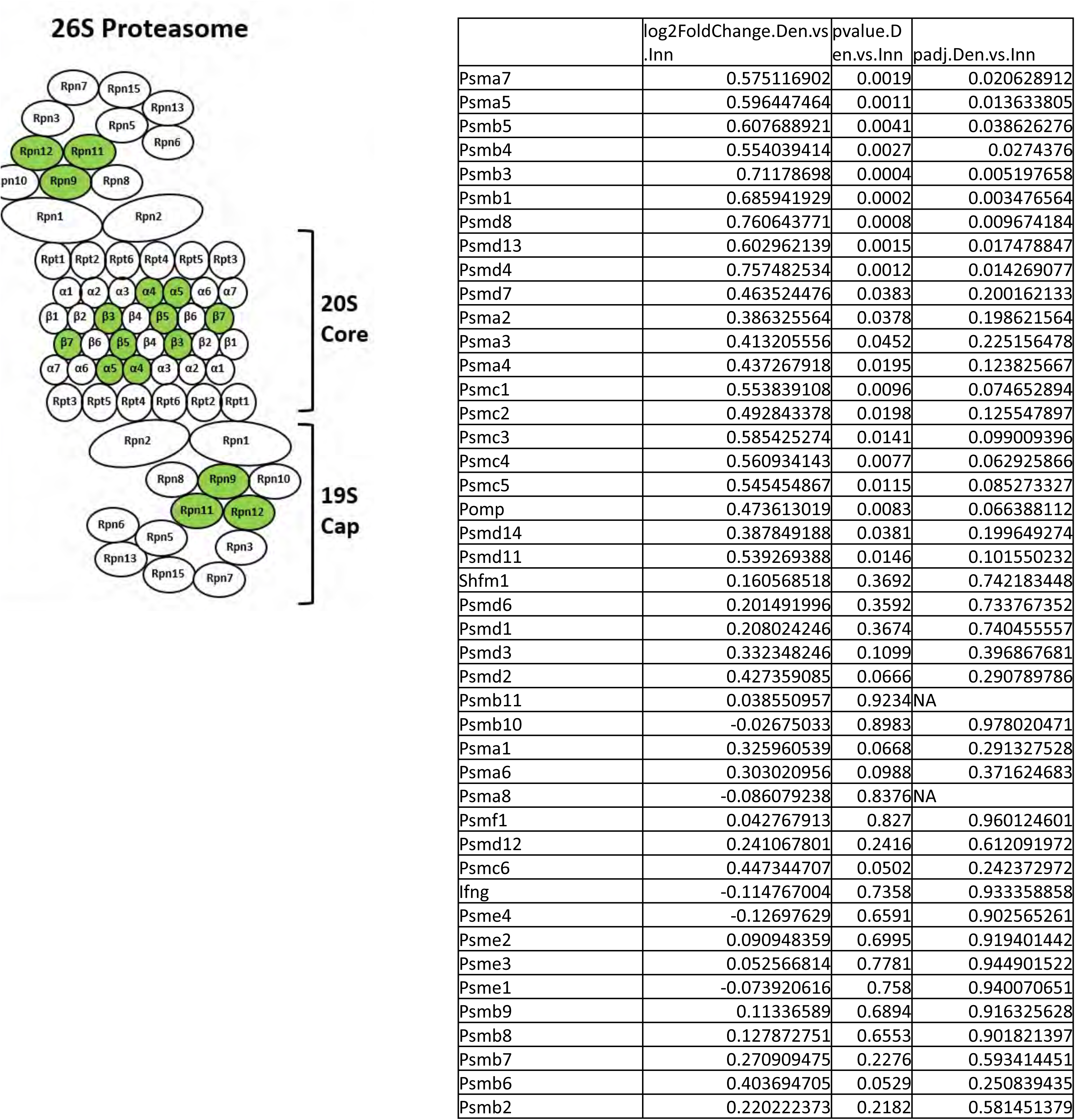
Expression of most proteasome genes return to basal levels in muscle at 14 d after denervation. To gain a broad view of transcriptional changes in the late phase of atrophy, 14 day denervated TA muscles were compared to innervated muscles by RNA-seq. Transcripts of several proteasome subunits and one proteasome chaperone were identified, and nine subunits were significantly induced in denervated compared to innervated muscles (induced subunits shown in green).

**Figure S2.**
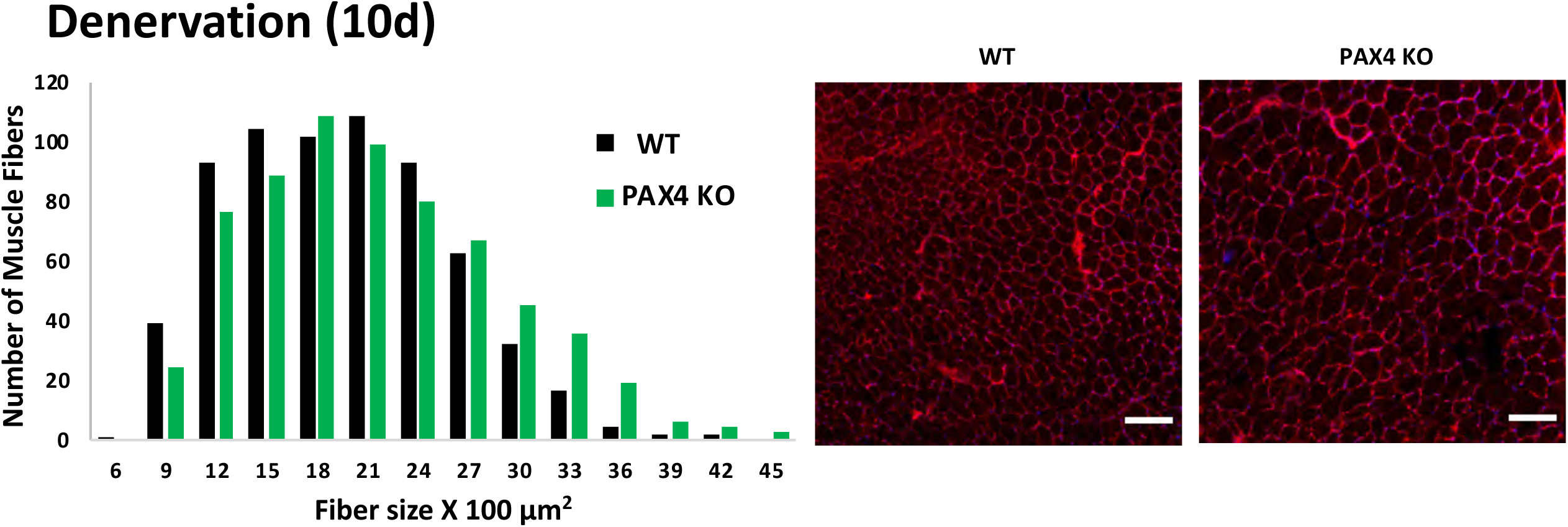
PAX4 deficiency attenuates muscle atrophy. Loss of PAX4 reduces fiber atrophy on denervation. Measurement of cross-sectional areas of 661 fibers from 10 d denervated muscles from WT *vs.* 661 fibers from PAX KO mice. N= 3 mice. Statistics is presented in table S1. Scale bar: 50 μm.

**Figure S3.**
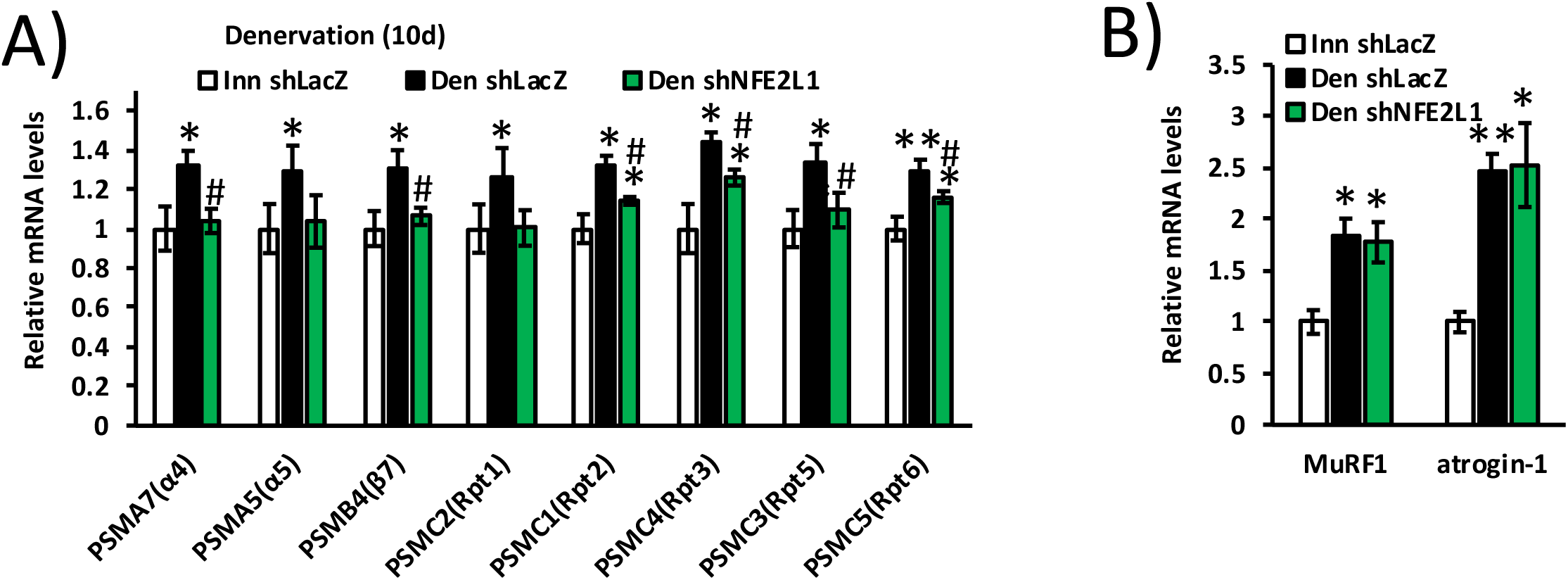
NRF-1^NFE2L1^ downregulation reduces the expression of some proteasome genes in denervated muscle. A-B) Expression of proteasome genes (A) and UPS components (B) was measured by RT-PCR analysis of mRNA preparations from innervated and 10 d denervated muscles expressing shNFE2L1 or shLacz. N=4. *, P < 0.05 *vs.* innervated shLacz; ** P < 0.001 *vs.* innervated shLacz; # P < 0.05 *vs.* denervated shLacz. Error bars represent SEM.

**Figure S4.**
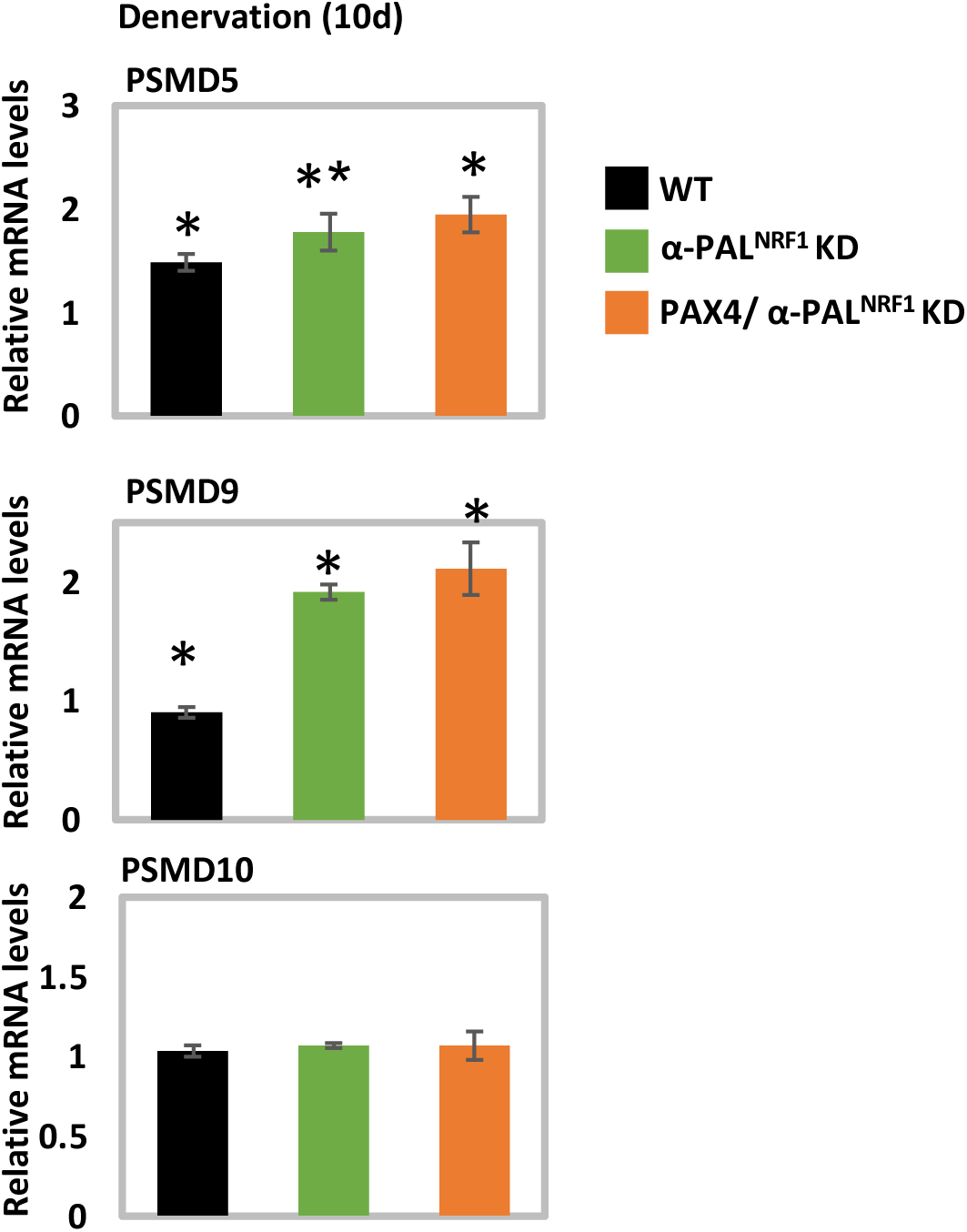
Neither PAX4 nor α-PAL^NRF-1^ controls the expression of the proteasome assembly chaperon genes PSMD5, PSMD9 or PSMD10. mRNA preparations from denervated (10d) muscles from WT, α-PAL^NRF-1^ KD, and PAX4/ α-PAL^NRF-1^ KD mice were analyzed by RT-PCR. N=4 mice. *, P < 0.05 and ** P < 0.001 *vs.* WT innervated controls. Error bars represent SEM.

**Table S1.**
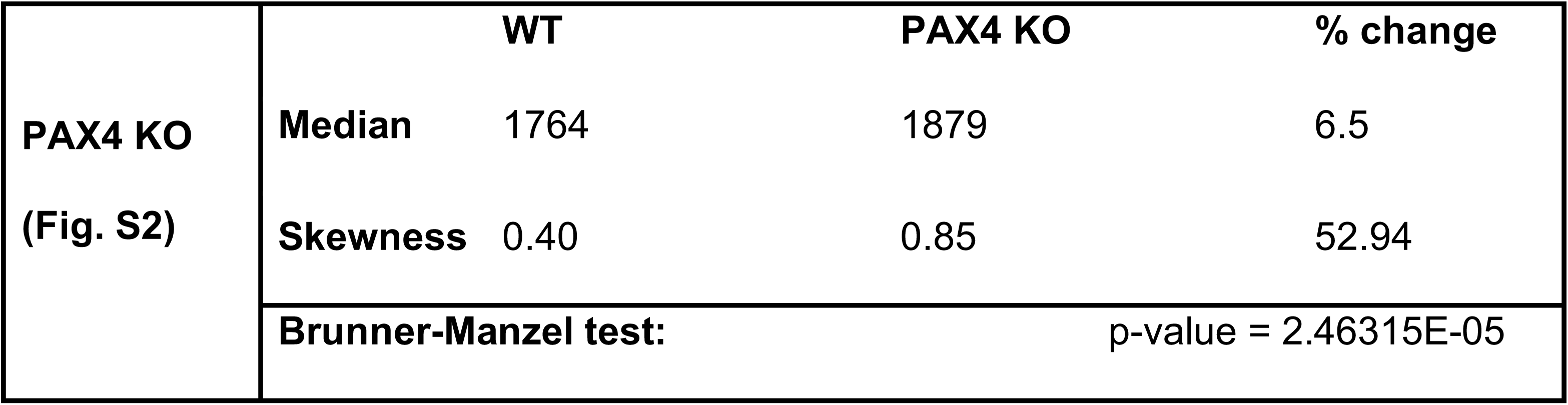
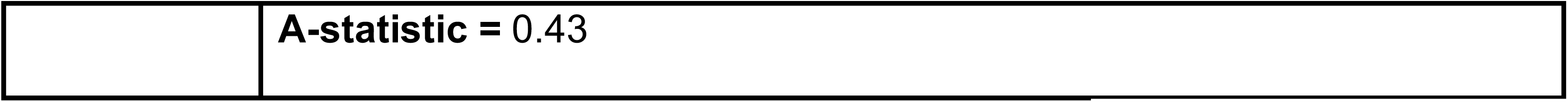
Summary statistics of fiber size analyses presented in Fig. S2 based on our recent methodology paper ^52^. With regard to A-statistics, if 0≤A<0.5 then dataset1 (WT) is stochastically less than dataset2 (PAX4 KO). The A-statistics is a direct measure of the fiber size effect ^52^, and it shows beneficial effect on cell size by the specific shRNAs. Such an effect can be simply missed by traditional measurements of median, average, and Student’s *t*-test.

|  |  | WT | PAX4 KO | % change |
| --- | --- | --- | --- | --- |
| **PAX4 KO**<br>**(Fig. S2)** | **Median** | 1764 | 1879 | 6.5 |
|  | **Skewness** | 0.40 | 0.85 | 52.94 |
|  | **Brunner-Manzel test:** | p-value = 2.46315E-05 |  |  |

|  |  |
| --- | --- |
|  | **A-statistic = 0.43** |

**Table S2.**
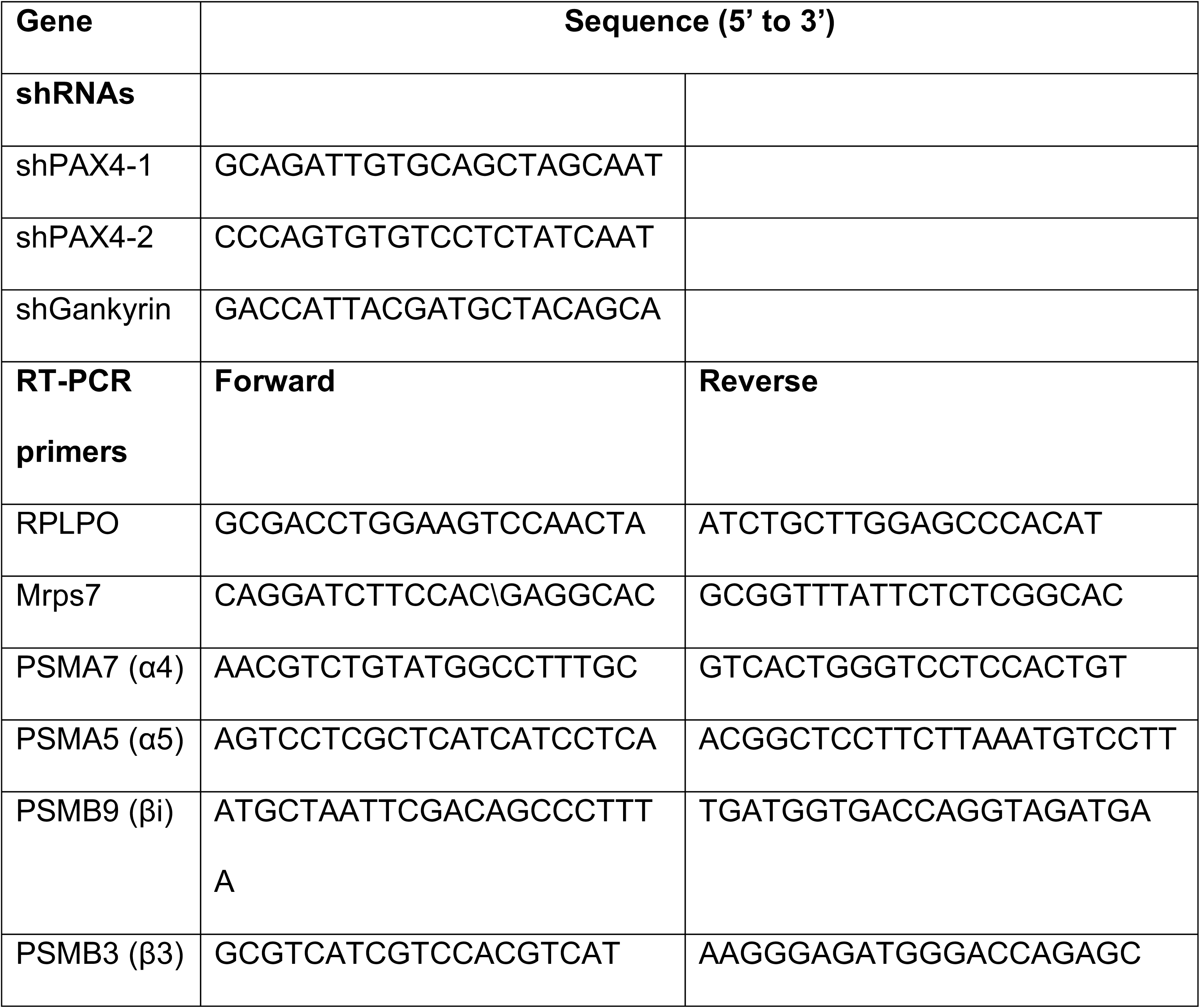
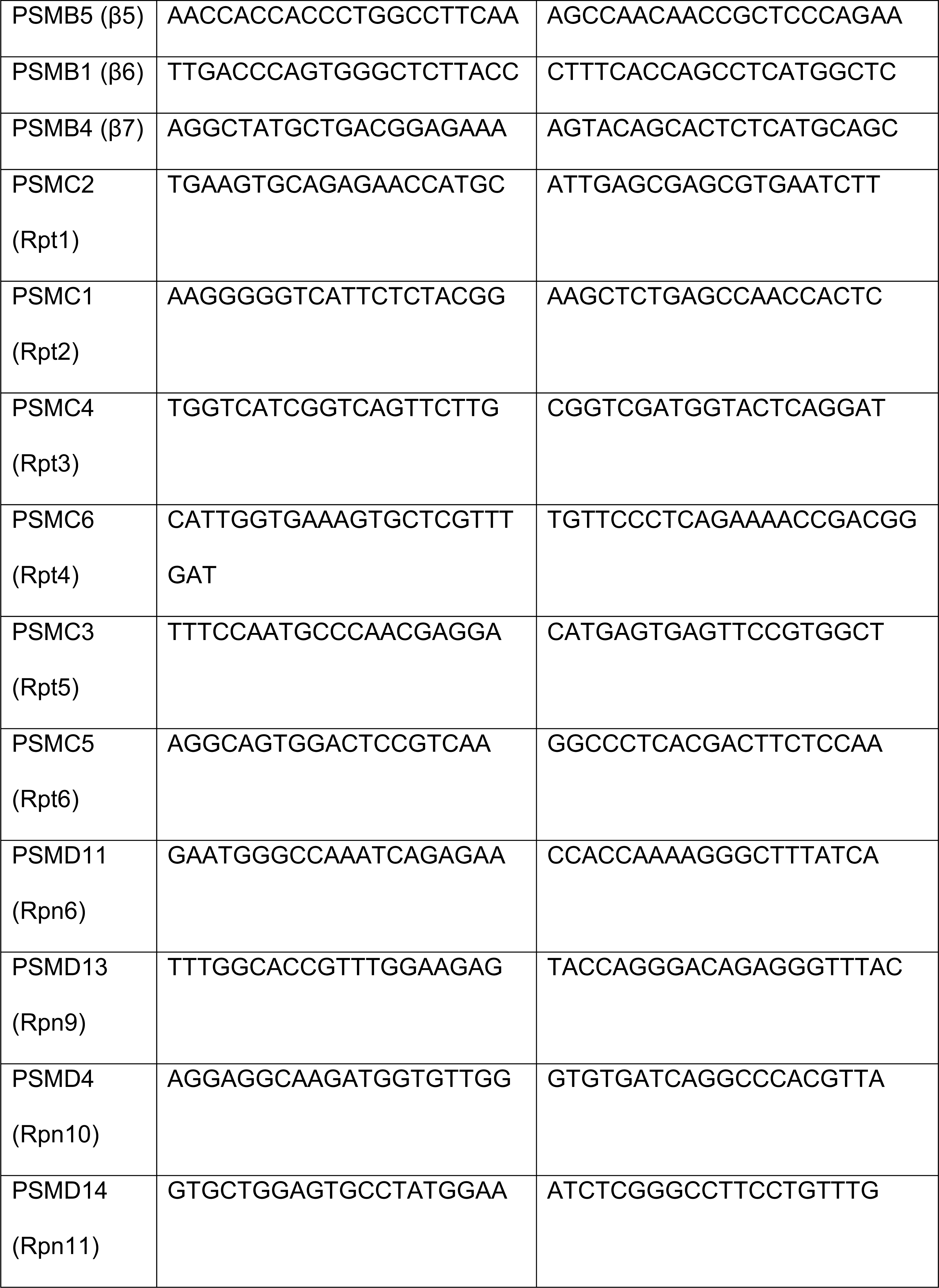
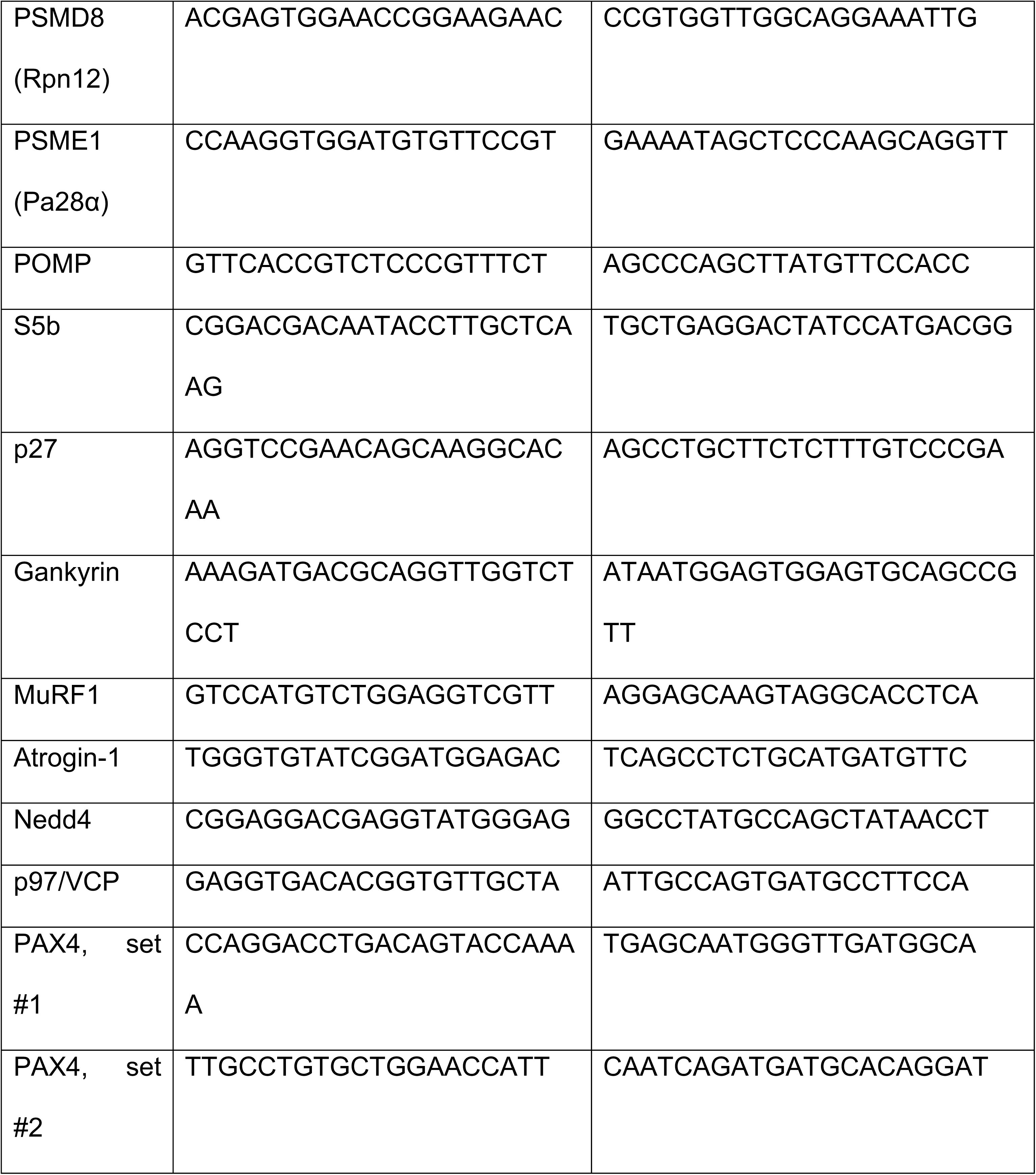
Primers used for RT-PCR

**Table S3.**
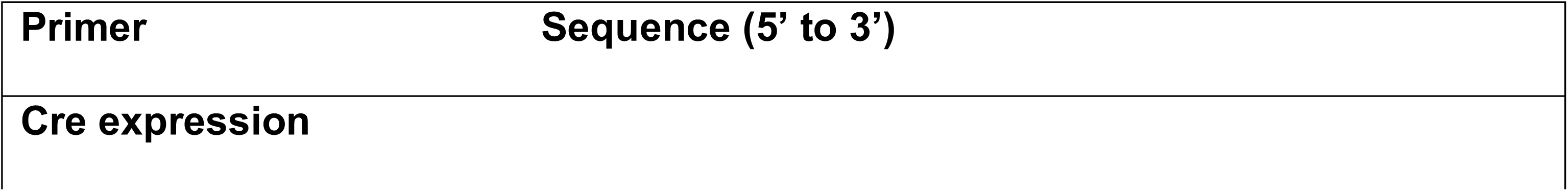
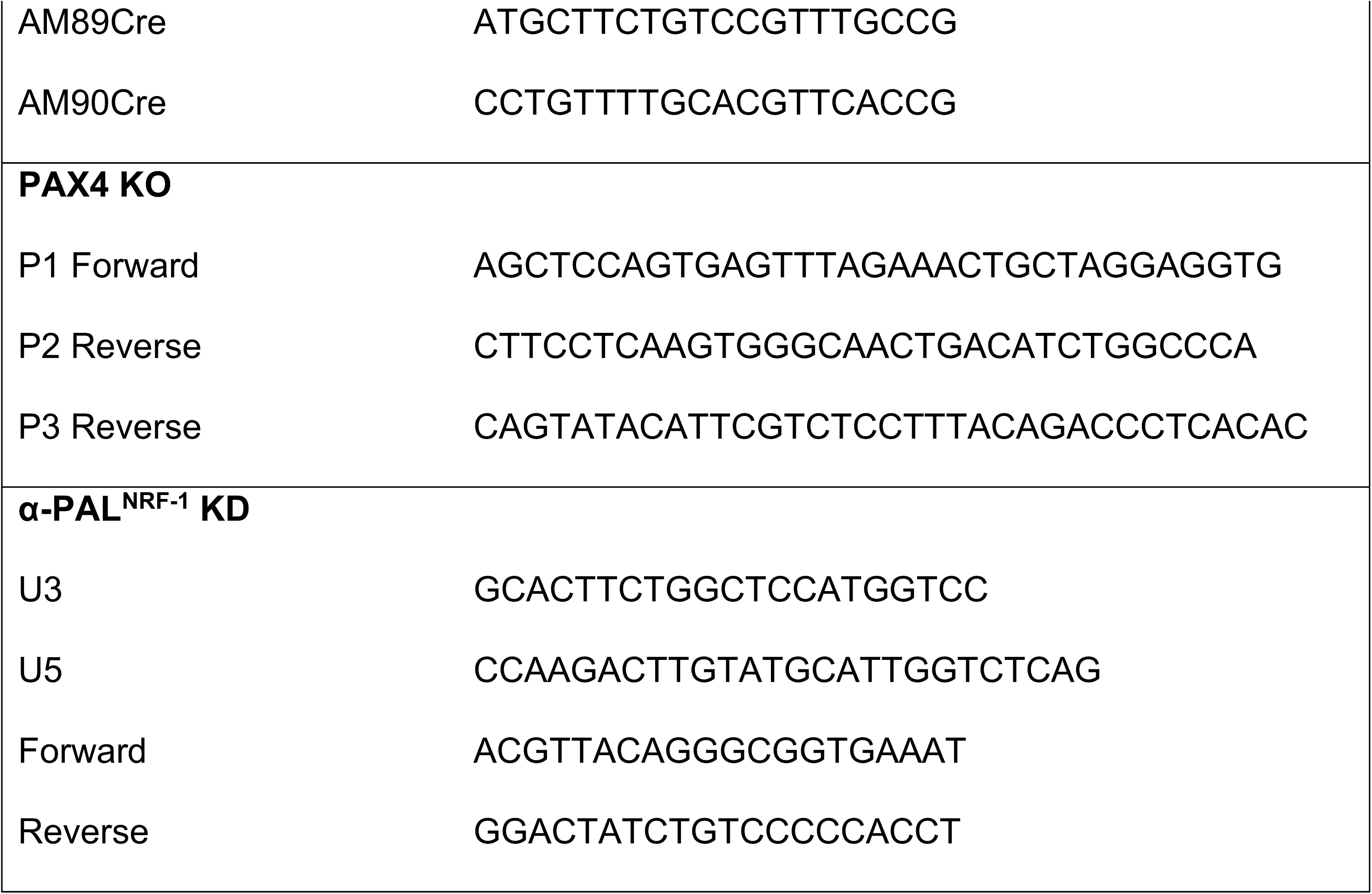
Primers used for Genotyping of PAX4 and NRF1 KO mice. Cre positive mice were identified using primers AM89Cre/AM90Cre; PAX4 floxed allele was identified with primers P1/P2; PAX4 KO was confirmed using primers P1/P3. α-PAL^NRF-1^ floxed allele was identified using primers U5 and U3; α-PAL^NRF-1^ KD was confirmed using α-PAL^NRF-1^ forward and reverse primers.

